# Pleiotropic mutation in a tendril TCP gene underlies the yield-enhancing *multiple-flowering* trait in summer squash (*Cucurbita pepo*)

**DOI:** 10.64898/2025.12.25.696487

**Authors:** Galil Tzuri, Adi Faigenboim-Doron, Harry S. Paris, Amit Gur

## Abstract

Crop yield is a focal point in plant breeding. Regulation of lateral budding through apical dominance was a central target of crop domestication, directly affecting crop production. The young fruits of *Cucurbita pepo*, summer squash, are produced on plants characterized by apical dominance and differentiation of a single flower bud per leaf axil. A single recessive mutation, *mf*, results in differentiation of more than one flower per leaf axil, thereby directly increasing production because of the continual day-to-day harvest of the summer-squash crop. Positional cloning of the *Cucurbita pepo mf* (*Cpmf*) gene denoted a frame-shift mutation in a TCP transcription factor, *Cp4.1LG13g07780*, as causative for the increase in axillary flowering. *Cpmf* is an ortholog of a tendril-development TCP gene in other cucurbits and, likewise, the recessive allele of *Cpmf* is associated with distorted tendril development. Gene function is context-dependent, and we propose that multiple-flowering is a unique pleiotropic attribute of mutation in a tendril-development gene of *C. pepo*. Characterization of a *C. pepo* collection confirmed a significant association of the *Cpmf* mutation with multiple-flowering, and showed that the mutant allele is absent in ancestral *C. pepo* and one of its two cultivated subspecies. The beneficial mutation occurred and was selected after the domestication of the other subspecies, during its cultivation for young fruit production. We demonstrate the discovery of a causative yield-increasing sequence variant and its practical utilization in breeding. Our findings provide a molecular target for creation of high-yielding, multiple-flowering summer-squash cultivars through marker-assisted breeding or precise genome editing.

**SIGNIFICANCE STATEMENT:** Summer squash is characterized by apical dominance and a single flower bud per leaf axil. A multiple-flowering trait, conferred by a single recessive gene, *mf*, significantly increases fruit yield by producing more flowers per leaf axil. We show that a mutation in a tendril-development TCP gene is causative for enhanced axillary flowering, and that the beneficial mutation occurred during the cultivation of summer squash. We demonstrate the successful implementation of *mf* for development of high-yielding squash cultivars.

## INTRODUCTION

Developmental architecture is a significant evolutionary attribute of plants. Common to the domestication of many plants is the transition of plant architecture toward apical dominance, expressed as suppression and regulation of lateral budding, and this transition is one of the phenomena defining the domestication syndrome (Hammer, 1984; Harlan, 1992; Heslop-Harrison and Schwarzacher, 2012; Meyer and Purugganan, 2013). Lateral budding is a major determinant of plant architecture as it involves the differentiation and subsequent development of branches, inflorescences, flowers, leaves and tendrils. Prime examples are maize (*Zea mays* L., Poaceae) and tomato (*Solanum lycopersicum* L., Solanaceae). In maize, the transition from its wild ancestor, teosinte, involved a dramatic reduction in lateral branching and an increase in yield derived from selection on the *tb1 (teosinte branched1)* gene (Doebley *et al*., 1997). In tomato, discovery of determinate growth triggered the development of processing tomato industry (Yeager, 1927). The causative *sp (self*-*pruning*) gene not only affects shoot determinacy but also regulates the transition of axillary meristems from vegetative to reproductive growth (Pnueli *et al*., 1998). Regulation of budding therefore, is a crucial yield-potential trait of crop plants grown for fruit or seed production.

Fruit or grain yield is strongly affected by resource allocation within the plant. In many cases, yield potential of a crop is not fulfilled due to reduced fruit setting regulated by limited internal resource availability. This is especially true of cucurbit crops. The fruits of the domesticated Cucurbitaceae are large and often contain several hundred seeds each, being strong sinks for plant assimilates. Developing cucurbit fruits place a strong demand on plant resources, thereby preventing the development of subsequent fruits and inhibiting further flowering and vegetative development. This inhibition has long been documented for melons (*Cucumis melo* L.) (Rosa, 1924; McGlasson and Pratt, 1963; Pratt *et al*., 1977) and winter squash (*Cucurbita maxima* Duchesne) (Bushnell, 1920; Zack and Loy, 1981; Loy, 2004). Even for cucumbers (*Cucumis sativus* L.), ‘first-fruit-inhibition’, was shown to be a limiting factor for higher yield (Schapendonk and Brouwer, 1984; Marcelis, 1993; Zhang *et al*., 2015; Shnaider *et al*., 2018).

Summer squash (*Cucurbita pepo* L.), another major cucurbit crop, are harvested and consumed as young fruits, 0–5 days past anthesis. As inhibition by developing cucurbit fruits sets in at around 6 days past anthesis and lasts for 10–14 days (Stephenson *et al*., 1988), timely continual removal of young fruits preempts inhibition of further flower, fruit, and vegetative development. As a result, in this crop, resource allocation is fundamentally different from that of the other major cucurbit crops because the continual removal of its young fruits allows for the continued, unfettered production of more leaves, flowers, and fruits, with the number of pistillate flowers produced becoming the main yield-defining component (El-Keblawy and Lovett-Doust, 1996).

*Cucurbita pepo* L. (Cucurbitaceae) is one of the most diverse and cosmopolitan of the cultivated cucurbits, encompassing hundreds of cultivars of summer squash, pumpkins, winter squash, and gourds (Paris, 2000). This species is grown in areas with temperate and sub-tropical climates the world over with a commercial impact that can be estimated at several billion dollars annually. Although the mature fruits, ≥40 days past anthesis, of pumpkins and squash are widely grown and appreciated for their flavor, nutritional value, and antioxidant content, and are also valued as ornamentals, it is the young, glossy fruits (0–5 days past anthesis), known as summer squash, that bestow most of the monetary value on this crop species.

*Cucurbita pepo* (2n□=□2x□=□40) is native to North America. It was first domesticated 10,000 years ago in central or southern Mexico and again 5,000 years ago in the eastern United States, resulting in two major cultivated lineages, which are taxonomically referred to as *C. pepo* subsp. *pepo* and *C. pepo* subsp. *ovifera* (L.) D.S. Decker (Smith, 2006). For thousands of years in North America, these two lineages evolved separately, in geographical isolation from each other. In spite of the many commonalities of growth and development, though, the two subspecies show some obvious and consistent differences between them (Paris *et al*., 2012). Among the traits that differentiate between the two subspecies is the ability or lack of ability to differentiate more than one flower bud in any given leaf axil on the main stem. Plants of subsp. *pepo* can generate only one flower bud per leaf axil but plants of subsp. *ovifera* can differentiate more than one. If two or more buds differentiate in an axil, they do not become visible to the naked eye simultaneously, but do so sequentially (Loy, 2004). The ability to differentiate more than one flower bud in a leaf axil, which has been termed *multiple flowering*, is Mendelianly inherited as a single recessive gene designated *mf* (**Figure 1a**) (Paris, 2018; Paris and Hanan, 2010).

**Figure 1:**
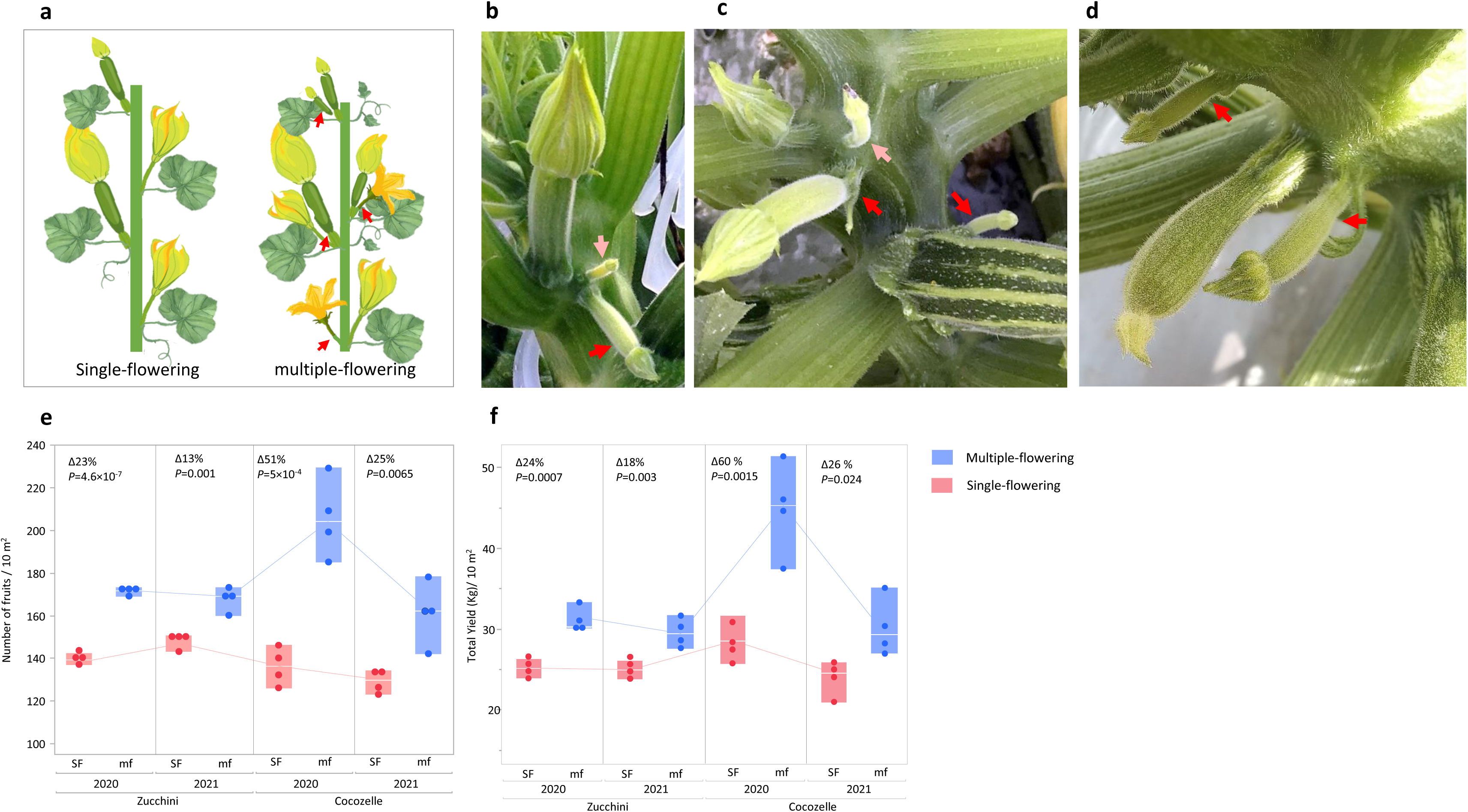
The multiple-flowering (*mf*) trait. (**a**) Schematic illustration of the plant architecture of squash with a normal single flower per leaf axil (SF) and the recessive *multiple-flowering* phenotype (mf). Red arrows indicate the secondary bud in leaf axils of the mf plant. (**b-d**) Photographs of multiple-flowering Cocozelle and Zucchini. Red and pink arrows indicate the secondary and tertiary buds in leaf axils, respectively. (**e**) Effect of the multiple-flowering trait on total yield in near-isogenic Cocozelle and Zucchini squash hybrids, genotype *mf/mf* (multiple-flowering) versus genotype *Mf/Mf* (single-flowering). After Paris and Gur (2022) (**f**) Effect of the multiple-flowering trait on number of marketable fruits in near-isogenic Cocozelle and Zucchini squash hybrids, genotype *mf/mf* (multiple-flowering) versus genotype *Mf/Mf* (single-flowering). After Paris and Gur (2022).

The edible-fruited cultivars, the pumpkins and squash, of the two cultivated subspecies have been sorted into eight morphotypes or cultivar-groups or, in short, Groups, based on fruit shape. Four of the Groups, Cocozelle, Pumpkin, Vegetable Marrow, and Zucchini, belong to subsp. *pepo* and the other four, Acorn, Crookneck, Scallop, and Straightneck, belong to subsp. *ovifera* (Paris, 2000). Two of the Groups, Pumpkin and Acorn, are grown mostly for production of their mature fruits, which are commonly referred to as pumpkins and winter squash. The other six Groups, though, are grown almost entirely for production of their young fruits, and these are known collectively as summer squash. Summer squash are an easy-to-grow, short-season but labor-intensive crop (Paris, 2008). Typically, summer squash fields are harvested every day or every other day. If not picked on time, the fruits continue to grow and become dull, tough, and unsalable. Theoretically, at least, the multiple-flowering trait of subsp. *ovifera* provides greater fruit yield potential for summer squash, because these young fruits are removed from the plant before becoming strong sinks for plant resources (Stephenson *et al*., 1988; El-Keblawy and Lovett-Doust, 1996). However, this trait was absent from *Cucurbita pepo* subsp. *pepo* until the introgression of the *mf* allele from a cultivar of the Crookneck Group of subsp. *ovifera* into subsp. *pepo* (Paris and Hanan, 2010).

The *mf* allele has since been introgressed by us into several inbreds of summer squash from both, the Cocozelle Group and the Zucchini Group of subsp. *pepo* (Paris, 2000) (**Figure 1b-d**), with the objective of determining whether the multiple-flowering trait could enhance yield in these two commercially highly valuable summer squash cultivar-groups. Into four of our inbreds, two Cocozelle and two Zucchini, we introgressed the trait to the sixth backcross generation (BC_6_), obtaining near-isogenic lines (NILs) with and without multiple-flowering, genotypes *mf/mf* and *Mf/Mf* (**Supplementary Figure S1**). We then crossed these near-isogenic inbreds to obtain two pairs of near-isogenic *mf/mf* and *Mf/Mf* hybrids, one pair for Cocozelle and one pair for Zucchini, and tested yield performance of these pairs of hybrids under field conditions. A significant yield-enhancing effect of the *mf* gene was shown in both backgrounds. For the Cocozelle, we observed as much as a 51% increase in fruit production with deployment of *mf* and, for the Zucchini, as much as a 23% increase (**Figure 1e, f**) (Paris and Gur, 2022).

Genomic research in *Cucurbita pepo* has become more effective since the release of a reference genome assembly of Zucchini in 2018 (Montero-Pau *et al*., 2018). This advancement was paralleled with construction of the Cucurbit Genomics Database (CuGenDB) (Zheng *et al*., 2019; Yu *et al*., 2023), jointly providing tools for genetic dissection of traits to the gene level in this crop species.

Given the demonstrated potential for markedly increased yields offered by *multiple-flowering* in subsp. *pepo*, our goal was to discover the molecular basis of the multiple-flowering trait. Herein, we describe the positional cloning of the *Cucurbita pepo mf (Cpmf)* gene and define a single-base insertion within a *TCP* transcription factor as the causative polymorphism for the increased axillary flowering. We show that this mutation is prevalent across the diversity of summer squash in *C. pepo* subsp. *ovifera* and is significantly associated with natural variation in the number of flowers per leaf axil, with signature for the evolution and selection of this yield-enhancing trait under domestication. *Cpmf* is the ortholog of the cucurbit tendril-development TCP gene in cucumber and melon (Wang *et al*., 2015; Mizuno *et al*., 2015), and we observed that it also affects tendril development in *C. pepo*, suggesting that multiple-flowering is a unique pleiotropic meristematic attribute of this mutation.

## RESULTS

### Whole-genome mapping of the multiple-flowering trait by BSA-Seq of nearly isogenic population

For genetic mapping of the multiple-flowering trait, we used the near-isogenic lines that were developed through 6 generations of backcrossing of the *mf* allele into single-flowering Zucchini and Cocozelle summer squash. The two BC_6_F_2_ populations were scored for single-or multiple-flowering phenotype. Analysis of both populations re-confirmed the recessive single-gene inheritance pattern of the multiple-flowering trait (**Figure 2a**). We selected 24 multiple-flowering and 15 single-flowering Zucchini segregants for bulk-segregant analysis by sequencing (BSA-Seq). Genome-wide allele frequency analyses supported the near-isogenic nature of the BC_6_F_2_ progenies, as more than 98% of the genome in both the multiple-flowering and single-flowering bulks coincided with the single-flowering recurrent Zucchini parent (TRF) allelic profile (**Figure 2b**). Only two narrow genomic regions, on chromosomes 7 and 13, displayed different patterns, with a significant signature for introgression from the multiple-flowering Crookneck donor parent (SET) (**Figure 2b**). However, while the chromosome 7 introgression was present in the expected random allele frequencies of 0.50 in both the multiple-flowering and the single-flowering bulks, and is therefore not related to the multiple-flowering trait, the chromosome 13 introgression displayed a significant ΔSNP-index signature, such that the multiple-flowering bulk displayed a high frequency of the SET allele (>0.80) and the single-flowering bulk displayed a frequency that matched the expected combination of 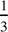 *Mf/Mf* and 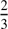 *Mf/mf* genotypes within the dominant phenotypic bulk in F_2_ (∼0.35 for the mf (SET) allele, **Figure 2c**). These results strongly pointed on a ∼900 Kb interval on chromosome 13 as the confidence interval for the *Cpmf* gene (**Figure 2c**).

**Figure 2:**
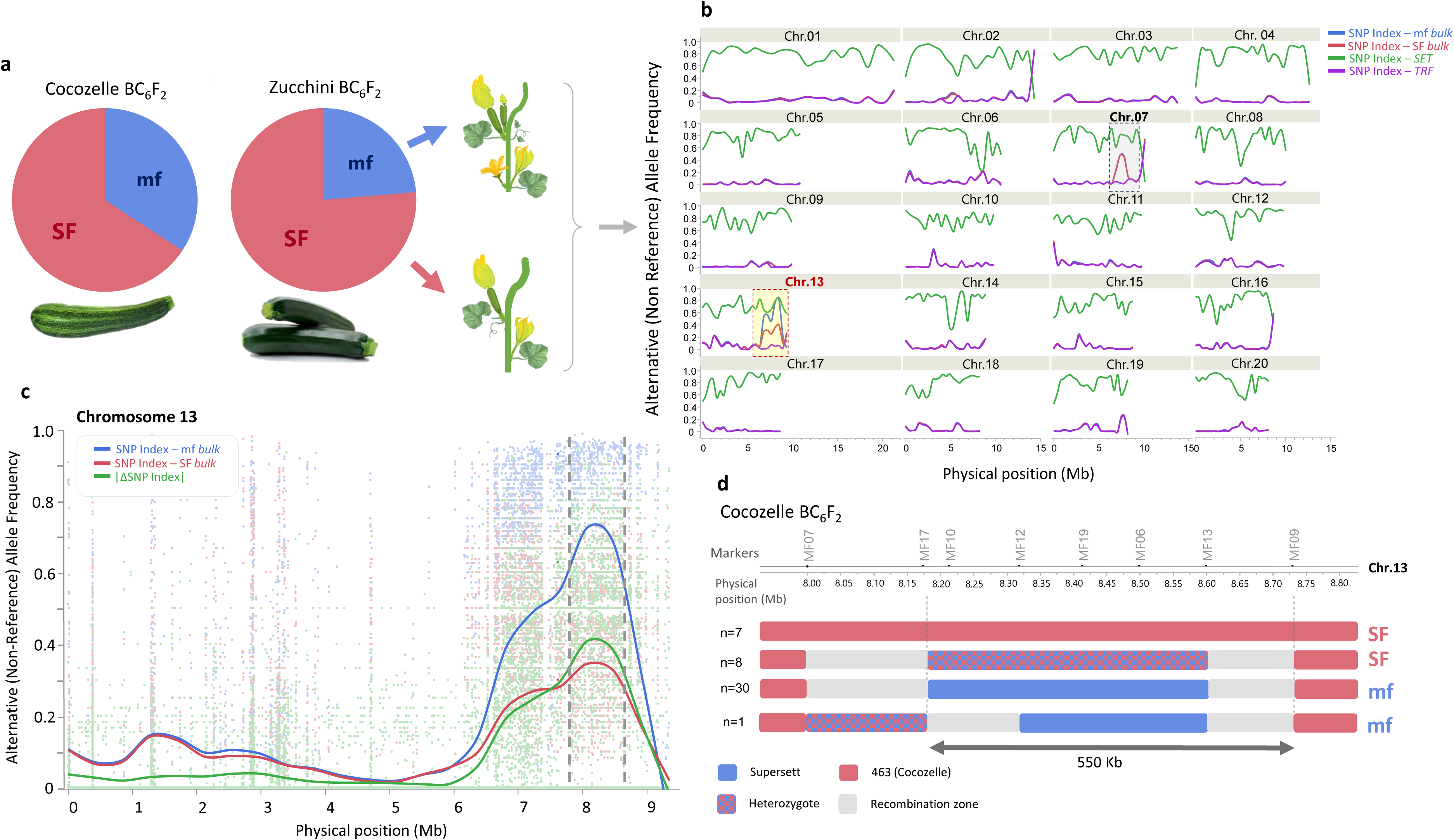
Whole-genome mapping of *multiple flowering*. (**a**) The phenotypic segregation of the single-flowering (SF) and multiple-flowering (mf) phenotypes in two BC_6_F_2_ populations introgressed with the Crookneck *mf* allele into Cocozelle and Zucchini backgrounds. The ∼3:1 SF: mf ratio is as expected for segregation of a single recessive gene. (**b**) The results of a bulk-sequencing analysis (BSA-Seq) whereby a significant association is detected on chromosome 13. Trend (running average) lines are presented for allele frequencies (SNP-index) in each of the bulks and the parents, Crookneck ‘Supersett’ (SET) and Zucchini ‘True French’ (TRF). (**c**) SNP-index analysis at the chromosome 13 *mf* region. Trend (running average) lines are presented for allele frequencies (SNP-index) in each of the bulks and the calculated ΔSNP-Index. Dashed vertical lines represent the trait confidence interval at 7.80 – 8.80 Mbp. (d) Genomic profile of the narrow chromosome 13 Crookneck ‘Supersett’ introgression in the BC_6_ of Cocozelle Accession 463 and BC_6_F_2_ segregants characterized using specific PCR markers (MF07, MF17, MF13 and MF09, **Table X**). At the left of each genotypic group the number (*n*) of BC_6_F_2_ plants is represented.

To validate the BSA-Seq results, we developed InDel Markers at the Chromosome 13 denoted interval and genotyped each of the individual plants that comprised the multiple-flowering and single-flowering bulks (**Supplementary Tables S2**, **S3**). This analysis confirmed the significant association with the chromosome 13 interval and allowed us, from eight recombinants that occurred in this region, to refine the trait interval to 823 Kb (8.152 Mb – 8.975 Mb) (**Supplementary Table S3**).

Subsequently, we analyzed 46 selected Cocozelle multiple-flowering and single-flowering plants and found that, in this genetic background, the initial donor introgression size in the chromosome 13 target region in the BC_6_ Accession 1951 was only 725 Kb (8.004 Mb – 8.729 Mb, **Figure 2d**), and that it perfectly co-segregated with flowering pattern (**Supplementary Table S3**). Of these, a single informative BC_6_F_2_ recombinant further narrowed the interval to 556 Kb (8.173 Mb – 8.729 Mb, **Figure 2d**).

### Fine-mapping of multiple-flowering to a single-gene resolution

To further narrow down the 556 Kb trait interval on chromosome 13, we genotyped, in three steps, a total of ∼1,200 BC_6_F_2_ seeds from the Zucchini population, for selection and analysis of recombinants within the target interval. We started with the flanking markers MF9 and MF17 and mapped the trait to a 286 Kb interval defined by markers MF6 (8.498 Mb) and MF10 (8.212 Mb) (**Figure 3a**). We then analyzed additional recombinants and zoomed in on a 95 Kb interval defined between markers MF12 (8.318 Mb) and MF19 (8.413 Mb). Analysis of 10 additional recombinants within this region allowed us to narrow down the mapping to a 4.5 Kb interval between markers MF29 (8.363 Mb) and MF26 (8.367 Mb) with a single annotated gene (**Figure 3b**). We then analyzed, with additional markers, four recombinants within this interval and narrowed the mapping to a 1.6 Kb interval between markers 7780_SNP#9 and marker MF31 (**Figure 3c**).

**Figure 3:**
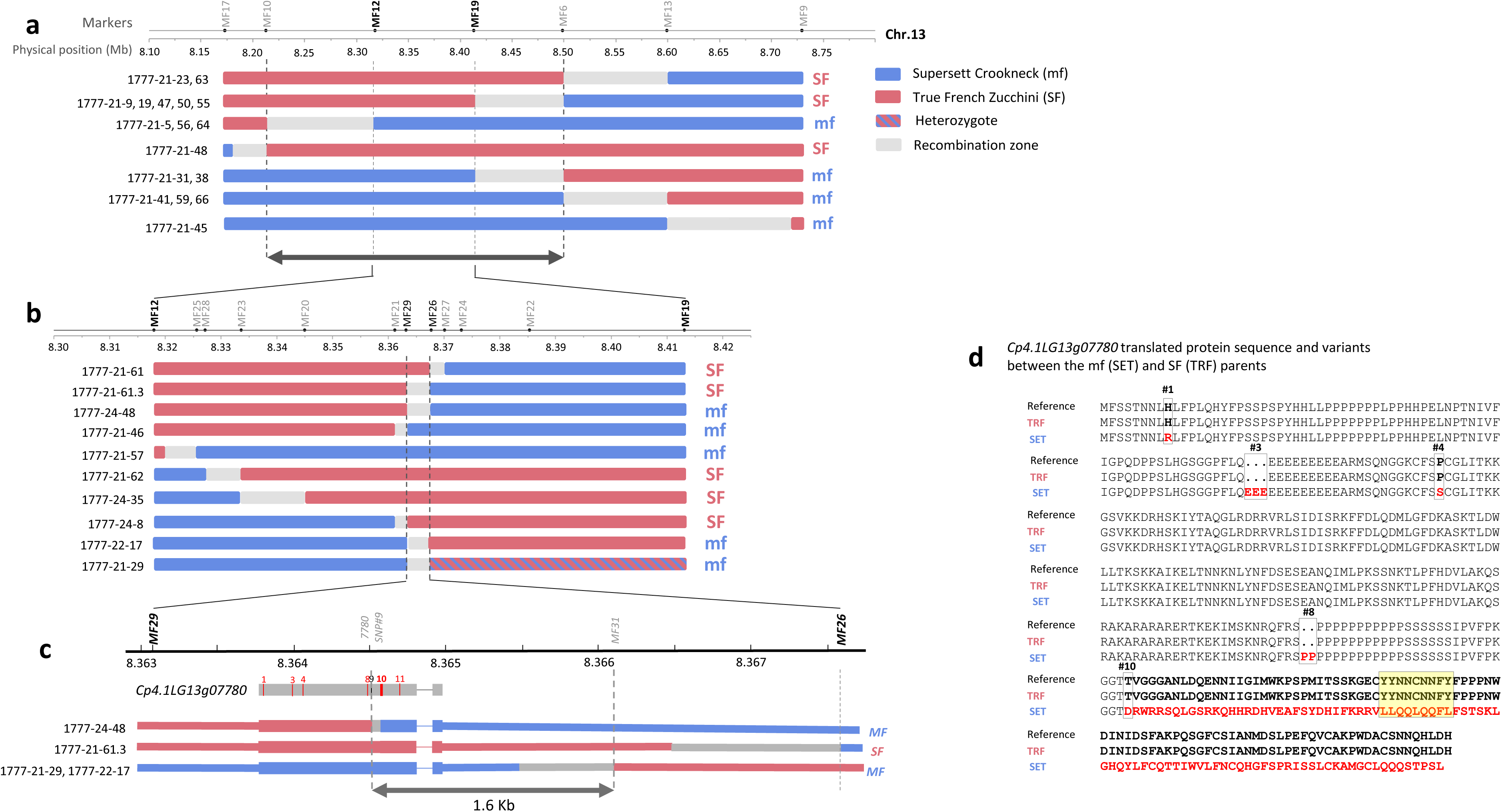
Positional cloning of the *Cpmf* gene. (**a**) Illustration of the BC_6_F_2:3_ recombinants between markers MF17 and MF9 used for the first round of substitution mapping to a ∼28Kb interval on *Cucurbita pepo* Chromosome 13 (*SF* = single flowering, *MF* = multiple flowering). (**b**) BC_6_F_2:3_ recombinants between markers MF12 and MF19 used for the second round of substitution mapping to a ∼6Kb interval with a single gene, *Cp4.1LG13g07780*, annotated as TCP Transcription-factor DICHOTOMA-like. (**c**) BC_6_F_2:3_ recombinants between markers MF29 and MF26 used for the third round of substitution mapping to a narrow 1,600 bp interval defining a single-base Insertion/Deletion (InDel) within the Cp4.1LG13g07780 gene (#10) as the most probable causative sequence variant for the multiple-flowering phenotype. The gene model is shown in its physical genomic coordinates. Red vertical numbered lines are non-synonymous polymorphisms between the SF (‘True French’ Zucchini) and MF (‘Supersett’ Crookneck) parents. (**d**) Cp4.1LG13g07780 protein sequence and variants between the multiple- and single-flowering parents. Polymorphism #10 is the single base-pair InDel causing the frame-shift and altered protein sequence (in red), including within a conserved Cucurbits-specific box (yellow rectangle).

### *Cpmf* is a TCP transcription factor

Only one gene, *Cp4.1LG13g07780*, annotated as a TCP (<u>T</u>B1, <u>C</u>YC, and <u>P</u>CF) transcription factor, is located within the 1.6 Kb interval defined by the fine-mapping (**Figure 3c**). The transcript structure of *Cpmf* was determined based on the *Cucurbita pepo* reference genome annotation (Montero-Pau *et al*., 2018) and using genomic and cDNA sequencing. The 1115-nucleotide transcript has two exons, and encodes a polypeptide of 339 amino acids with the highly conserved TCP domain in the first exon. The minimal mapping interval restricted the multiple-flowering causative variant to the sequence starting towards the end of the coding region of the TCP gene and extending to the 3’UTR region (**Figure 3c**). Comparative analysis of the genomic and cDNA sequences of the parents, TRF and SET, revealed 18 polymorphic sites within this gene, 11 within exon and 7 in intronic sequence (**Supplementary Figure S2 and Supplementary Table S4**). Six of these polymorphisms are non-synonymous changes that impact the translated protein sequence (**Figure 3d, and Supplementary Table S4**). SNPs #1, #4 and #11 are single amino acid (AA) substitutions. Polymorphism #3 is a three AA insertion/deletion (InDel), #8 is a two AA InDel, and polymorphism #10 is a single base InDel with the most prominent effect on the protein sequence. This InDel induces a frameshift that modifies the sequence of the last 91 amino acids at the C-terminus of the protein (**Figure 3d**), and was therefore the strongest candidate as the causative variant for the multiple-flowering trait. The BC_6_F_3_ recombinant 1777-24-48 defined the left border of the trait interval at SNP#9 at 8,364,514 bp (**Figure 3c**) and therefore also positionally confirmed InDel#10 (at 8,364,572 bp) as the most probable causative candidate polymorphism. SNP#11 is embedded within the shifted protein sequence as it is located 36 AA downstream to InDel#10 and is therefore a less probable candidate. Modification of the last 91 AA induced by InDel#10 is predicted to affect the TCP transcription factor activity as this region includes modification of a conserved motif reported in cucumber (Wang *et al*., 2015) (**Figure 4**).

**Figure 4:**
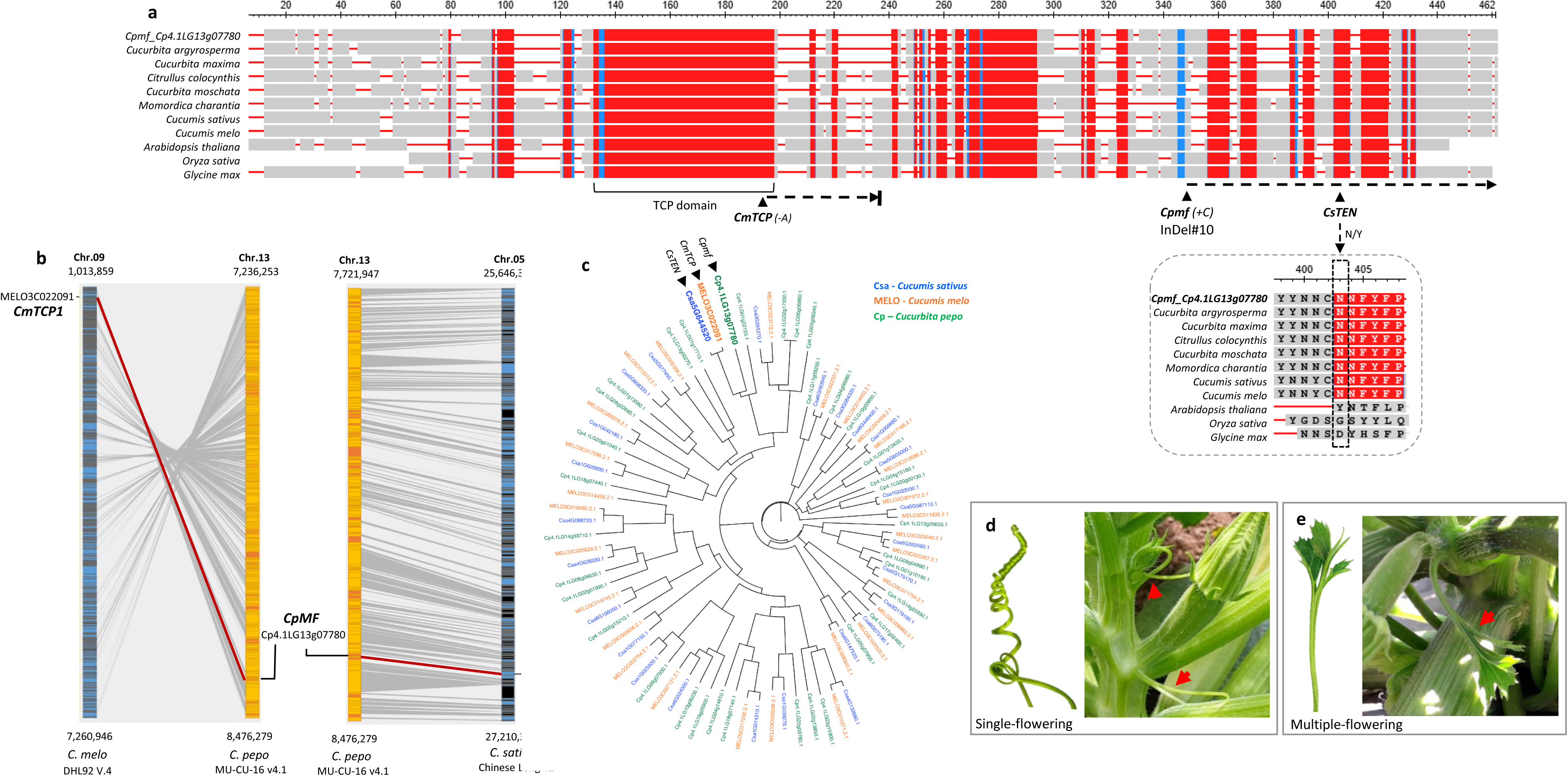
*Cpmf* is a TCP gene, the ortholog of the cucurbit tendril gene. (**a**) TCP Protein sequence alignment across 11 plant species. Red boxes represent highly conserved sequences within the Cucurbitaceae. Gray boxes are aligned homologous sequences. Red horizontal lines are gaps. The TCP motif is labeled at the bottom. The positions of causative polymorphism sites are labeled for *Cucurbita pepo* (*Cpmf*), *Cucumis melo* (*CmTCP1*) and *Cucumis sativus* (*CsTEN*), including zoom in on the conserved cucurbits motif described by Wang *et al*. (2015). (**b**) Syntenic blocks of the *Cpmf* orthologs in *C. melo* and *C. sativus*. (**c**) Phylogenetic tree of 91 TCP proteins from *C. pepo*, *C. melo* and *C. sativus* (from Wang *et al*., 2025) including *Cpmf*, *CmTCP1* and *CsTEN*. (**d**) Normal coiled tendril in the ‘True French’ Zucchini (single flowering, *Mf/Mf*). (**e**) Distorted leaf-like tendril (red arrow) in the ‘True French’ Zucchini (multiple flowering, *mf*/*mf*, near-isogenic, Accession 1777).

### *Cpmf* is the ortholog of a tendril-development gene of cucumber and melon and the mutant allele shares a common loss of tendril-identity phenotype

Protein sequence comparisons and phylogeny, followed by analysis of synteny, suggested that the *Cpmf* gene (*Cp4.1LG13g07780*) is the ortholog of *CmTCP1* (*MELO3C022091*) and *CsTEN* (*Csa5G644520*) genes in melon and cucumber, respectively (**Figure 4a-c**). Both genes were previously shown to be specifically associated with regulation of tendril development in these cucurbit species (Mizuno *et al*., 2015; Wang *et al*., 2015). Indeed, detailed phenotypic comparison between the multiple-flowering and single-flowering near-isogenic squash lines revealed, in addition to the variation in the number of flowers per leaf axil, also a difference in tendrils development and morphology. While single-flowering segregants showed normal curled cylindrical tendrils (**Figure 4d**), the multiple-flowering segregants tend to develop distorted, split or leaf-like tendrils (**Figure 4e, Supplementary Figure S3).** Other than these phenotypic differences, near-isogenic lines and hybrids differing in the multiple-flowering introgression were phenotypically similar in fruit and plant characteristics (**Supplementary Figure S4**). These defined phenotypic effects are found in accordance with the specific *Cpmf* expression pattern in the stem at leaf axils (SLA) and tendrils (TEN) (**Figure 5**), which is in line with the tendril-specific expression reported for *CmTCP1* (*MELO3C022091*) and *CsTEN* (*Csa5G644520*) genes (Wang *et al*., 2015; Mizuno *et al*., 2015) (**Supplementary Figure S5)**, and providing further support for their orthology.

**Figure 5:**
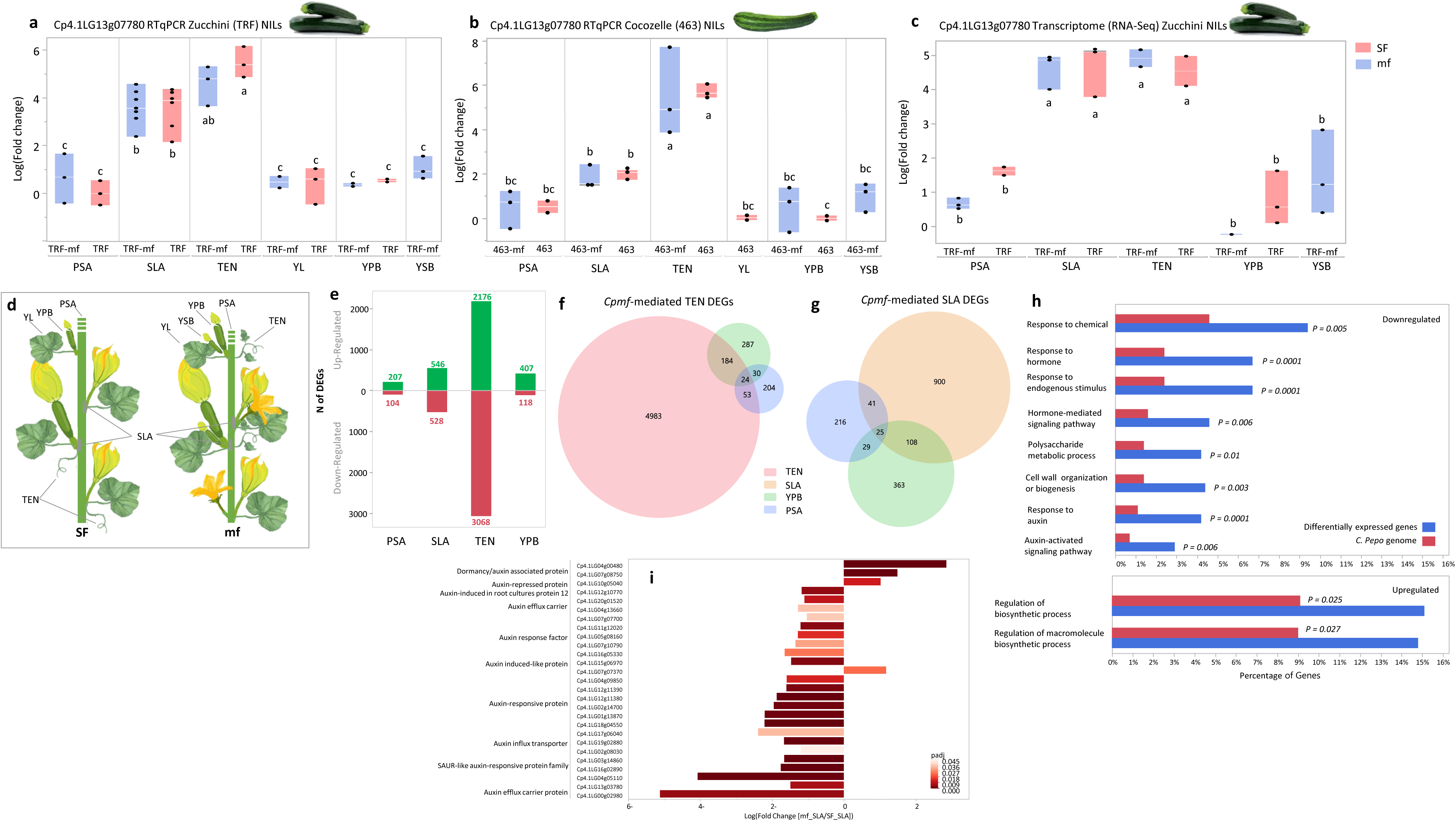
Expression of the *Cpmf* and downstream genes across tissues in near-isogenic lines (NILs) in Zucchini and Cocozelle backgrounds. (**a**) Comparison of expression of *Cp4.1LG13g07780* (by qRT-PCR) between multiple-flowering (TRF-mf) and single-flowering (TRF) Zucchini NILs across six tissues. Each point represents a biological replication collected from 2 plants. PSA = primary shoot apex, SLA = shoot at leaf axil, TEN = tendril, YL = young leaf, YPB = young primary bud, YSB = young secondary bud (only in TRF-mf). Log (fold-change) values labelled with the same lower-case letter are statistically non-significant at *P* < 0.05. (**b**) Comparison of expression of *Cp4.1LG13g07780* (by qRT-PCR) between multiple-flowering (463-mf) and single-flowering (463) Cocozelle NILs across six tissues. Each point represents a biological replication collected from 2 plants. Abbreviations along *x*-axis as in (a), with YSB only in 463-mf. Log (fold-change) values labelled with the same lower-case letter are statistically non-significant at *P* < 0.05. (**c**) Comparison of expression of *Cp4.1LG13g07780* (by RNA-Seq) between multiple-flowering (TRF-mf) and single-flowering (TRF) Zucchini NILs across five tissues. Abbreviations along *y*-axis as in (a). Log (fold-change) values labelled with the same lower-case letter are statistically non-significant at *P* < 0.05. (**d**) Schematic representations of sampling sites on single-flowering (SF) and multiple-flowering (mf) plants. Abbreviations as in (a) and (b). (**e**) Up- and down-regulated genes (green and red bars, respectively) between multiple- and single-flowering NILs in the BC_6_F_2_ population across different tissues. PSA = Primary shoot apex, SLA = Stem at Leaf Axil, TEN = Tendril, YPB = Young Primary Bud. Numbers above and below each bar represent the number of differentially expressed genes (DEGs) in each category. (**f**) Tendril-specific DEGs. Venn diagram of DEGs between multiple- and single-flowering NILs in the BC_6_F_2_ Zucchini population across 3 tissues: Tendrils (TEN), Young fruits (YPB) and primary shoot apex (PSA). Circle sizes are proportional. The number in each area represents the number of DEGs. (**g**) Stem at leaf-axil (SLA)-specific DEGs. Venn diagram of DEGs between multiple- and single-flowering NILs in the BC_6_F_2_ population across 3 tissues: Stem at leaf axil (SLA), Young fruits (YPB) and primary shoot apex (PSA). Circle sizes are proportional. The number in each area represents the number of DEGs. (**h**) Gene ontology (GO) enrichment analysis of SLA-specific differentially expressed genes (DEGs). (**i**) Differential expression of 27 auxin-related genes.

### *Cpmf* is expressed in tendrils and in the stem at leaf axils and is not differential between multiple-flowering and single-flowering NILs

We characterized the expression profile of *Cp4.1LG13g07780* in six different tissues in the single-flowering and multiple-flowering NILs in both the Zucchini and Cocozelle backgrounds (**Figure 5a-d**). Expression pattern was very similar in both backgrounds and indicated that *Cp4.1LG13g07780* is not expressed in young leaves (YL), primary shoot apex (PSA) and young primary and secondary buds (YPB and YSB). In both backgrounds, the *Cpmf* gene is expressed in the stem at the leaf axils (SLA) and the strongest expression was found in tendrils (TEN). This analysis also showed that in both, the Zucchini and Cocozelle backgrounds, there is no significant difference in the expression of *Cp4.1LG13g07780* between the multiple-flowering and single-flowering NILs across the different tissues, suggesting that the multiple-flowering phenotype is not a result of differential expression of the TCP gene.

### Mutation in *Cpmf* is associated with downregulation of auxin-related genes at leaf axils

To gain further insight on the transcriptomic effect of allelic variation at *Cpmf*, we performed RNA-Seq on multiple tissues and compared between the multiple-flowering and single-flowering NILs. Thousands of genes were differentially expressed between the NILs, and majority of the differentially expressed genes (DEGs) were found in tendrils (TEN, 5,244 DEGs) and to a lesser extent in the stem at leaf axil tissue (SLA, 1,074 DEGs) (**Figure 5e**). Substantially smaller numbers of DEGs were found in young fruits and apical meristems (YPB, 525 DEGs and PSA, 311 DEGs). These results coincide with the specific expression and action of *Cpmf* in SLA and TEN tissues where the phenotypic impact of this gene is observed. To focus on the expression patterns in these two *Cpmf*-specific tissues, we substituted the DEGs detected in YPB and PSA, both of which are tissues where *Cpmf* was not expressed, from TEN and SLA DEGs, resulting in 4,983 TEN-specific DEGs (**Figure 5f**) and 900 SLA-specific DEGs (**Figure 5g, Supplementary Table S5**). Gene ontology (GO) enrichment analysis on the SLA-specific up- and down-regulated genes support the central role of this TCP gene in axillary meristematic activity. Sixty-three of the 418 up-regulated genes are classified as involved in the general regulation of biosynthetic process (**Figure 5h**), reflecting a significant 65% enrichment of genes in this process. The most prominent GO enrichment was found for down-regulated genes related to hormone signaling pathways, and more specifically to auxin signaling (4-fold enrichment, *P*<0.0001, **Figure 5h**). Among the SLA-specific DEGs, we found 27 genes annotated as auxin-related, and 25 of them displayed a significant 2 to 16-fold lower expression in the multiple flowering plants (**Figure 5i, Supplementary Table S6**). Regulation of apical dominance and axillary budding through auxin has been described in other plant species (Shen *et al*., 2019; Takeda *et al*., 2003; Aguilar-Martínez *et al*., 2007).

### Phylogeny and expression profile of the TCP gene family in Cucurbita pepo

In consensus with (Wang *et al*., 2025), 36 TCP genes were identified in the reference Zucchini genome (MU_-_CU_-_16 V4.1) (Montero-Pau *et al*., 2018) and they are distributed across 14 of the 20 *Cucurbita pepo* chromosomes. These TCPs are classified into 4 defined clades and *Cpmf* is located with its melon and cucumber orthologs — *CmTCP1* and *CsTEN* — on the same branch in Clade IIC that corresponds with the TB1/CYC1 subgroup (**Figure 6a**). Two-way hierarchical clustering of expression of these TCPs across different tissues in the *Cpmf* NILs revealed a good correlation between expression profile clustering and the phylogenetic clades (**Figure 6b-c**). Expression cluster #2 includes *Cpmf* and 3 other genes that display a common SLA and TEN specific expression pattern, and all belong to Clade IIC. *Cp4.1LG01g22120* is the closest paralog of *Cpmf* and also the most similar in its expression pattern, suggesting possible redundant or related functions of these two genes (**Figure 6b**). Another interesting group is the 3 TCPs in expression cluster #3 that is phylogenetically classified as Clade IIB and displays tendril-differential expression between the *mf* NILs, such that they are upregulated under the multiple-flowering allele of *Cpmf* (**Figure 6b**). This suggests that these TCPs are interacting with *Cpmf* and most likely are also involved in regulation of tendril development.

**Figure 6:**
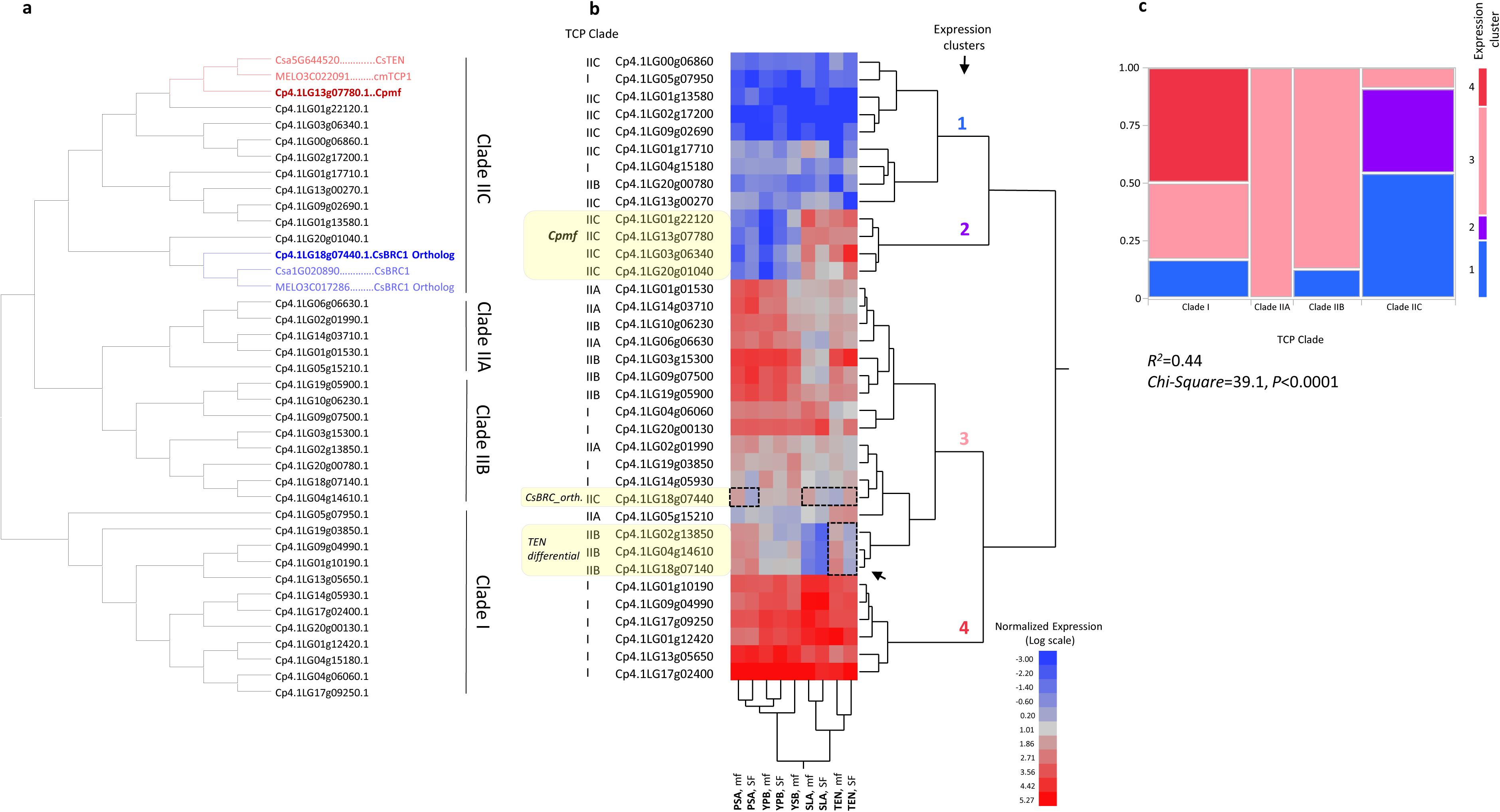
Phylogenetics and expression profiles of *Cucurbita pepo* TCPs. (**a**) Phylogenetic tree of 36 *Cucurbita pepo* TCP protein sequences. (**b**) Two-way hierarchical clustering of expression of 36 TCPs across 5 different plant tissues in the BC_6_F_2_ mf NILs. PSA = primary shoot apex, YPB = young primary bud, YSB = young secondary bud (only in TRF-mf), SLA = shoot at leaf axil, TEN = tendril, mf = multiple-flowering, SF = single-flowering. (**c**) Mosaic plot and chi-square analysis for the relation between expression clusters and TCP phylogenetic clades. Cell colors correspond to expression cluster numbering

### Number of flowers per leaf axil is a quantitative trait and significantly associated with allelic variation in the *Cpmf* gene

To expand our perspective on the axillary flowering trait across *Cucurbita pepo* diversity, we analyzed a diverse core set of 50 accessions that represents three subspecies and the eight edible-fruited cultivar-groups (Hereafter, *Core50*). We observed substantial variation for growth habit and plant architecture, and we quantified it by tallying variation in differentiation of lateral organs at leaf axils and measuring internode length, which are the main growth habit descriptors. Significant heritable variation was found for all 6 traits (**Figure 7a, Supplementary Figures S6-S11**). Correlation analysis (**Figure 7b**) showed that in consensus with the difference between vine and bush growth habits, internode length (InL) is positively correlated with number of side branches per leaf axil (nBr) and, interestingly, also positively correlated with the number of tendrils (nTEN). The number of female flowers (nFF) per leaf axil was negatively correlated with nTEN, nBr and InL across this panel (**Figure 7b**). Principle component analysis (PCA) using these 6 growth habit traits (**Figure 7c**) displayed the architectural differences between the two cultivated subspecies and highlight the significant developmental differences between the multiple-flowering Crookneck parent (SET) and the two single-flowering Zucchini (TRF) and Cocozelle (463) parents. While the Zucchini and Cocozelle near-isogenic segregating populations were scored categorically as multiple-flowering or single-flowering (**Figure 2a**), detailed analysis of the diverse panel revealed the quantitative nature of axillary flowering pattern with a range of 0.50 to 4 flowers per leaf axil (**Figure 7d**). Analysis of variance confirmed a very significant genetic effect for this trait (*H^2^* = 0.87, **Figure 7d**) and a significant difference in axillary flowering among the four cultivar-groups of subsp. *ovifera* (**Figure 7e**, *R^2^* = 0.68).

**Figure 7:**
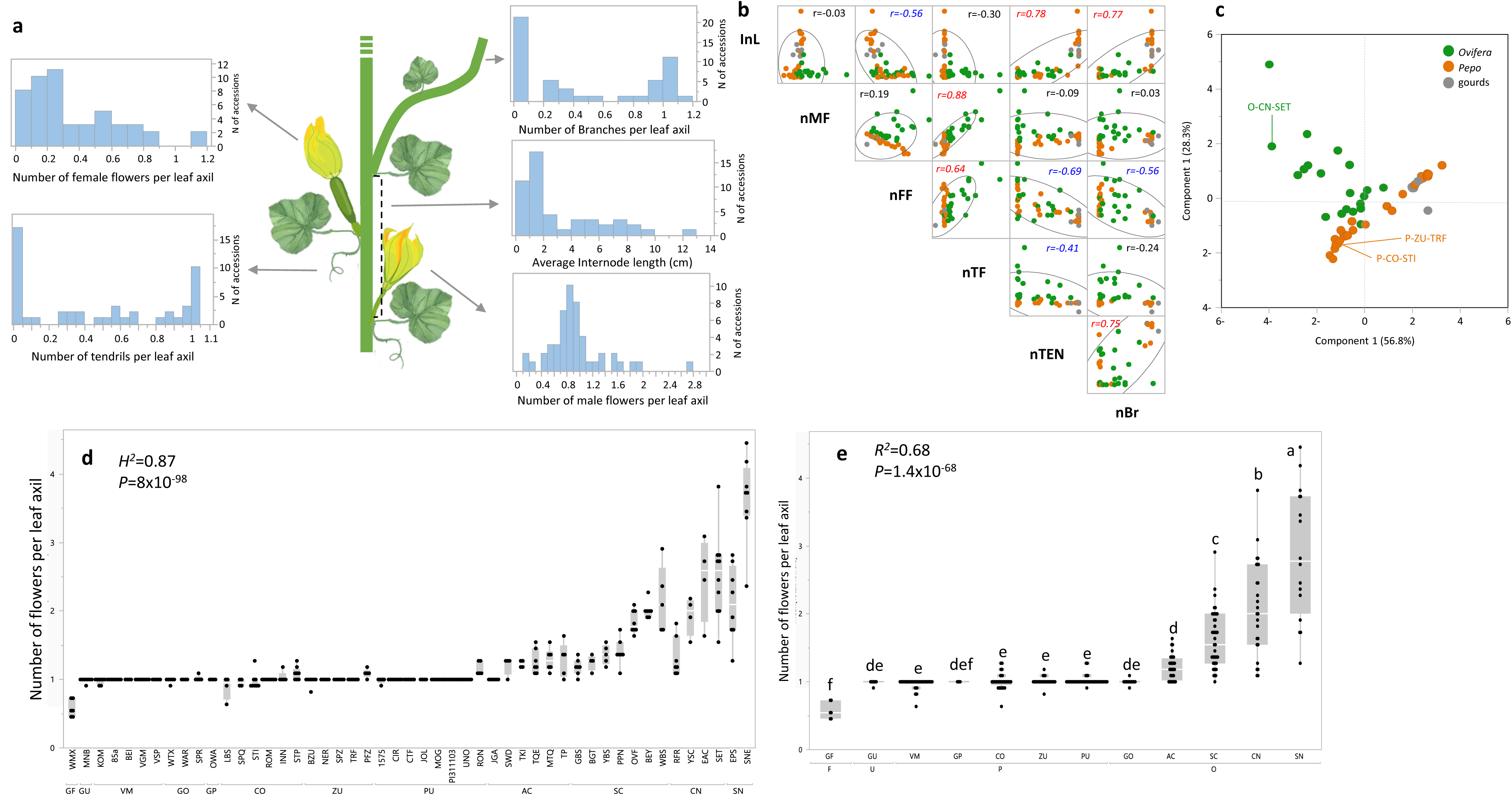
Variation in plant architecture and flowering pattern across diversity in *Cucurbita pepo*. (**a**) Distributions of plant architecture properties across 50 *Cucurbita pepo* accessions. (**b**) Pairwise correlation matrix between 6 plant architecture-related traits. Each point represents the mean of 11 leaf axils per plant and 4–8 plants per accession. Dot colors correspond with subspecific classification: green for *Ovifera*, orange for *Pepo*, and gray dots represent gourds. InL = internode length, nMF = number of male flowers, nFF = number of female flowers, nTF = total number of flowers, nTEN = number of tendrils, nBr = number of brunches. (**c**) Principal components analysis (PCA) of 6 plant architecture-related traits. Dot colors correspond with subspecific classification: green for *Ovifera*, orange for *Pepo*, and gray dots represent gourds. Parents of the near-isogenic populations are labeled. (**d**) Comparison of number of flowers per leaf axil across 50 *C. pepo* accessions. Each point is the average of 11 leaf axils per plant, from the 10^th^ through the 20^th^ on the main stem. Abbreviations for Groups and gourds are: GF = Gourd Fraterna, GU = Gourd Unclassified, VM = Vegetable Marrow, GP = Gourd Pepo, CO = Cocozelle, ZU = Zucchini, PU = Pumpkin, GO = Gourd Ovifera, AC = Acorn, SC = Scallop, CN = Crookneck, SN = Straightneck. (**e**) Comparison of number of flowers per leaf axil across the 8 cultivar-groups and 4 gourd taxa. Means labelled with a common letter are statistically non-significant at *P* < 0.05.

Hierarchical clustering of the *Core50* set based on genome-wide InDel and SSR markers (**Supplementary Table S7, S8**) is consistent with the subspecies differentiation and with genetic differences among cultivar-groups (**Figure 8a**) as also previously described in more detail (Gong *et al*., 2012; Paris *et al*., 2015). The *Cpmf* gene was sequenced across the *Core50* set and allelic variation was scored across all the polymorphic sites (**Supplementary Table S9**). InDel#10 within the *Cpmf* gene displays differences in allele frequencies between the *pepo* and *ovifera* subspecies, with the single-base insertion (+C), multiple-flowering allele, present only within subsp. *ovifera* (**Figure 8a**). While there is linkage disequilibrium (LD) between the adjacent polymorphisms within the *Cpmf* gene, InDel#10 showed the most significant association with the number of flowers per leaf axil across the diverse core panel (*R^2^* = 0.61, *P* = 3.5×10^-11^, **Figure 8b-c** and **Supplementary Table S10**), providing another support for this site as causative for the flowering variation.

**Figure 8:**
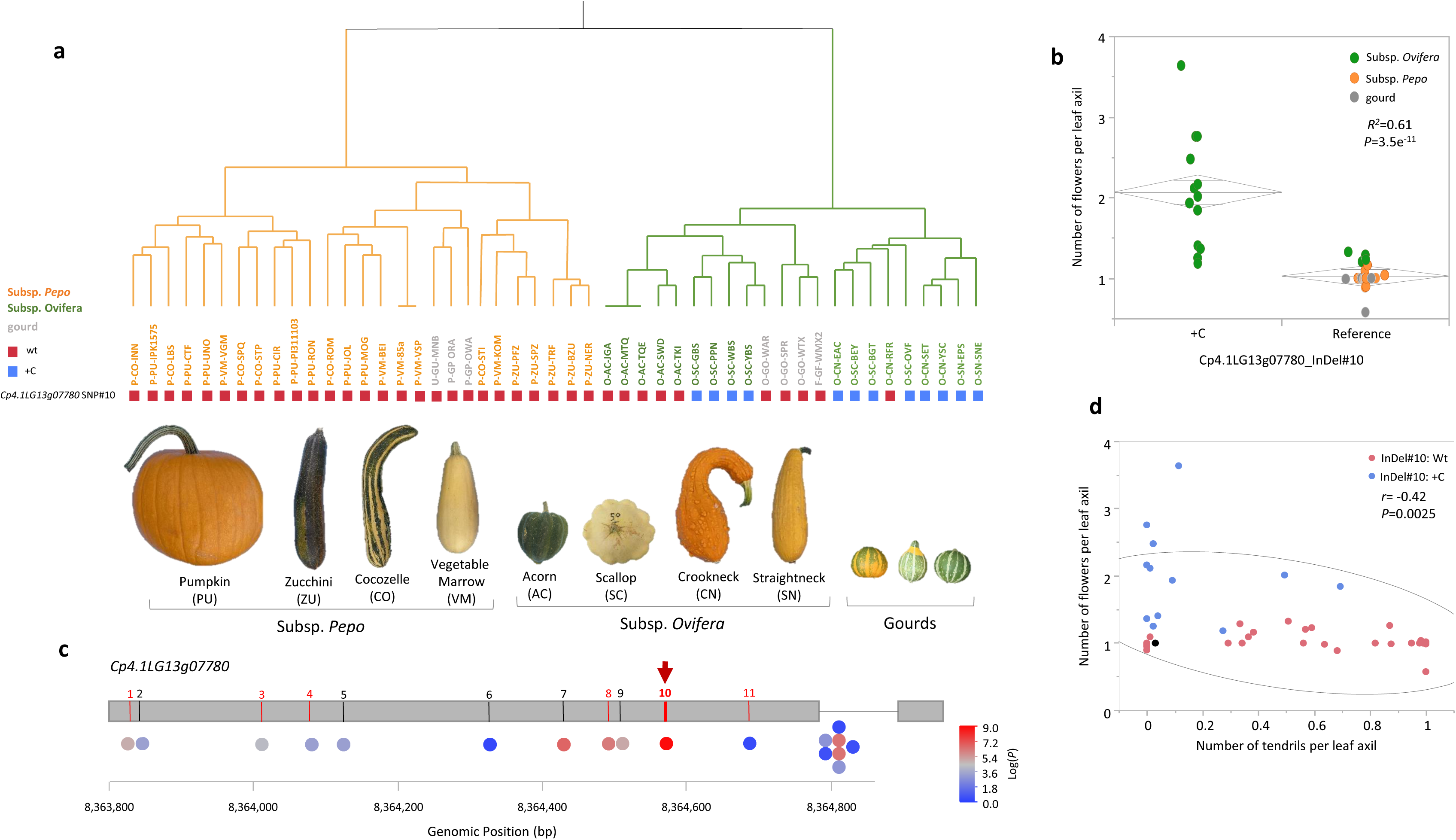
InDel#10 in the *Cpmf* gene is associated with the number of flowers and abundance of tendrils across a diverse *Cucurbita pepo* collection. (a) Hierarchical clustering of 49 core accessions (‘Supersett’ is represented once) based on genome-wide InDel markers variation (24 polymorphic sites **Supplementary Tables S7** and **S8**). Abbreviated accession names are colored according to sub-species, green for *Ovifera* and orange for *Pepo,* except that all gourds are colored in gray. Colored rectangles bellow the accession names represent the allelic variation at *Cp4.1LG13g07780* InDel#10. Blue is the *mf* (multiple-flowering) allele (+C insertion). Red is the wild-type *Mf* (single-flowering) allele. (**b**) Association of *Cp4.1LG13g07780* InDel#10 with number of flowers per leaf axil across 50 accessions. (**c**) Associations of 18 polymorphic sites, plotted along the *Cpmf* gene, with the number of flowers per leaf axil, as calculated across the core 50 *C. pepo* diverse panel. Polymorphic site # are denoted above the gene plot and corresponds to **Supplementary Tables S4.** Red and black colors represent non-synonymous and synonymous polymorphism, respectively. Red arrow is pointing on InDel#10. (**d**) Correlation between number of tendrils per leaf axil and number of flowers per leaf axil across the 50 accessions. Dot colors correspond with the genotype at *Cp4.1LG13g07780* InDel#10, blue is the *mf* allele (+C insertion) and red is the wild-type *Mf* allele.

Alongside the 43 edible-fruited accessions of subspecies *pepo* and *ovifera*, the *Core50* set included also 7 gourd accessions (labelled F-GF-WMX2, O-GO-SPR, O-GO-WAR, O-GO-WTX, P-GP-ORA, P-GP-OWA, U-GU-MNB), all of which displayed the single-flowering phenotype (Groups GP, GO, GU, **Figure 7e** and **Supplementary Table S1**). Interestingly, regardless of subspecific affiliation and regardless of whether they were collected in the wild or cultivated, these gourd accessions were monomorphic for the single-flowering genotype only at the InDel#10 site within *Cpmf* (**Figure 8a-b** and **Supplementary Table S9)**. Furthermore, a focused look on the allelic distribution of InDel#10 within subsp. *ovifera* reveals that of its four edible-fruited cultivar-groups, only the Acorn (AC) is monomorphic for the single-flowering allele (**Figure 8a**) and, indeed, this group displays a significantly lower number of flowers per leaf axil compared to the other three cultivar-groups within ssp. *ovifera* (**Figure 7e**). The Acorn Group cultivars are grown for consumption of their ripe fruits (winter squash).

The other three cultivar-groups of subsp. *ovifera*, Crookneck, Scallop, and Straightneck, are grown for consumption of their young fruits (summer squash). All but one of their accessions had the *mf* genotype at the InDel#10 site. The one exception was ‘Rugosa Friulana’ (RFR), a recently rediscovered heirloom from Italy, defined by its fruit characteristics and genotypic profile as a Crookneck (**Figure 8a**, **Supplementary Figure S12**). This accession has reduced axillary flowering which corresponds with its single-flowering genotype at the *Cpmf* causative site (**Figures 7d, 8a** and **Supplementary Table S9)**. Interestingly, this accession differs from all other Crookneck squash known to us in other traits, including stem and developmental fruit coloration. Possibly, this collection of unusual traits are the result of contamination in the ancestry of this old cultivar.

Collectively, these results further support the causality of *Cpmf* from InDel#10 mutation and provide evidence that this mutation along with the multiple-flowering phenotype most likely arose and was selected during the cultivation of subsp. *ovifera*. A significant negative correlation (*r* = −0.42, *P* = 0.0025) was also found across this panel between the number of flowers per leaf axil and presence of tendrils (**Figure 8d**), supporting the involvement of the *Cpmf* gene in regulating both traits.

## DISCUSSION

### The multiple-flowering trait is conferred by a pleiotropic mutation in the cucurbit tendril gene

Through positional cloning approach, we showed here that the multiple-flowering trait in *Cucurbita pepo* is a result of a single base InDel mutation in a TCP transcription factor gene (**Figures 2 and 3**). TCP proteins (TCPs) are plant-specific transcription factors that have a central role in the evolution and developmental control of plant form. TCPs are involved in diverse growth-related processes such as leaf development, branching, floral organ morphogenesis and hormone signaling (Martín-Trillo and Cubas, 2010). These proteins contain a highly conserved domain, the TCP domain, defined by the first identified members of the family: *TB1* (teosinte branched1), *CYC* (cycloidea), and *PCF1* and 2 (Cubas *et al*., 1999).

In cucurbits, specific TCP genes are key players underlying variation in tendril development and morphology (Sousa-Baena *et al*., 2018). Genetic analysis of the ‘Chiba Tendril-Less’ (ctl) melon (*Cucumis melo* L., Reticulatus Group), a mutant that completely lacks tendrils and develops lateral shoots instead, revealed that the tendril identity is determined by the gene *CTL*, which corresponds to *CmTCP1*, a melon transcription factor belonging to the TCP group of proteins (Mizuno *et al*., 2015). A tendril-less line of *Cucumis sativus* (CG9192) has also been identified in which tendrils are replaced by an organ with branch identity. This mutation in cucumber was shown to be conferred by the recessive allele of *CsTEN*, a TCP gene orthologous to the melon *CmTCP1* (Wang *et al*., 2015).

Using three layers of evidence (sequence homology, genomic synteny and parallel expression pattern, **Figures 4, 5**), we show here that *Cpmf* is the ortholog of both, *CmTCP1* and *CsTEN*, and belongs to the subgroup TB1/CYC1 of the Class II TCP gene family. TB1/CYC1 genes maintained their *TEOSINTE BRANCHED1-like* (*TB1-like*) role across different taxa, as they negatively regulate axillary bud outgrowth in monocots and eudicots (Dhaka *et al*., 2017). Accordingly, we found that the recessive *mf/mf* genotypes display abnormal tendril development in squash (**Figure 4d-e** and **Supplementary Figure S3**). In consensus with their effect on tendril development, both *CmTCP1* and *CsTEN* genes are expressed almost exclusively in tendrils (Mizuno *et al*., 2015; Wang *et al*., 2015). We found that the strongest expression of *Cpmf* is indeed in tendrils and that the expression level is not differential between *Mf* and *mf* genotypes (**Figure 4**), supporting the altered TCP protein sequence (and functionality) as the causative molecular modification.

As fundamental biological pathways are evolutionarily conserved across taxa, orthologous genes may affect variation in the same traits. However, it is not unlikely that functional differences arise between orthologues during speciation (Gabaldón and Koonin, 2013). Our genetic investigation suggests that a novel pleiotropic function evolved for the cucurbit tendril-specific TCP gene in *Cucurbita pepo* — increased axillary flowering (the multiple-flowering attribute) — supported by the expression of the *Cpmf* gene also in the stem at leaf axils and not exclusively in tendrils (**Figure 5**).

In cucumber, axillary lateral branching was shown to be regulated by a different TCP gene (*CsBRC1*) (Shen *et al*., 2019). The *Cucurbita pepo* ortholog of *CsBRC1* (*Cp4.1LG*18*g07440*, based on homology and segmental synteny, **Figure 6a, Supplementary Figure S13**) has a different expression profile compared to *Cpmf* (**Figure 6b**) and could be a potential candidate as a regulator of lateral branching, which is an important domestication trait in *C. pepo*. Interestingly, *Cp4.1LG*18*g07440* is differentially expressed under the different *Cpmf* alleles. It is up-regulated under the multiple-flowering allele in the primary shoot apex and in the stem at the leaf axil, and down-regulated in tendrils (**Figure 6b**). These results suggest some level of interaction between these two TCP genes.

### Evolution under cultivation of the multiple-flowering trait and its utilization for yield enhancement in summer squash

One of the iconic domestication genes in crop plants is *Teosinte Branched1* (*TB1*), which in domesticated maize suppresses the growth of axillary buds, enhancing apical dominance compared to its wild ancestor, teosinte (Doebley *et al*., 1997). A quarter-century ago, the TCP transcription factor gene family was defined (Cubas *et al*., 1999) and multiple homologous TCP genes were shown to cause parallel morphologies in other plant species, including *TB1* homologs in rice, wheat, and barley that were shown to regulate lateral branching and inflorescence architecture (Zhang *et al*., 2019; Ramsay *et al*., 2011; Dixon *et al*., 2018; Takeda *et al*., 2003). In maize, the domestication selection was directed to higher expression of *TB1,* which acts as a negative regulator of axillary budding (Doebley *et al*., 1997). Interestingly, we found that the selection that acted on *Cpmf*, a *Cucurbita pepo* homolog of *TB1*, was in the opposite direction, towards loss-of-function that promoted enhanced axillary flowering. We show here that the evolution of the multiple-flowering attribute was independent of apical dominance in *C. pepo*. The transition from the ancestral vine growth habit towards bush architecture and apical dominance is expressed as shorter internodes and reduced lateral branching, but is not significantly associated with variation in axillary flowering (**Supplementary Figure S14a-c**). Accordingly, the causative site *Cpmf-InDel#10* is not associated with axillary branching (**Supplementary Figure S14d)**. These results imply that different genes, still to be discovered, are regulating axillary branching in *C. pepo* and that *Cpmf* is specific to tendril and flower axillary meristems (**Figures 5a-c, 6b**). A similar independent regulation pattern is evident in cucumber where different TCP family members are involved in regulation of axillary tendril development (Wang *et al*., 2015) and lateral branching (Shen *et al*., 2019).

Wild plants of *Cucurbita pepo* are procumbent or climbing, highly branched vines bearing small, 3–8 cm diameter, spherical, globular, oval, or pyriform, bitter and toughly fibrous but smooth-rinded, prettily striped gourds, which likely attracted the attention of early hunter-gatherers (Nee, 1990). All seven of the *C. pepo* gourd accessions that we studied were single-flowering, regardless of subspecific affiliation (**Figure 8a**). Indeed, they were also monomorphic for the single-flowering allele at InDel#10 within the *Cpmf* gene (**Figure 8a-b, Supplementary Table S9**). The gourd accession Wild Arkansas (O-GO-WAR) is a representative of *C. pepo* subsp. *ovifera* var. *ozarkana* D.S. Decker, the wild taxon considered to be the direct ancestor of the cultivated taxon subsp. *ovifera* var. *ovifera* (Decker-Walters *et al*., 1993). The taxon of wild gourds, var. *ozarkana*, had widespread use by native peoples in what is now the eastern half of the United States by 8000 years ago (Smith, 2006). These gourds began to be domesticated as early as 5000 years ago (Cowan, 1997; Smith, 2006) via the ruderal pathway, benefitting from and adapting to changes in the environment resulting from disturbances caused by human activity (Fuller *et al*., 2025), which provided favorable landscapes for its growth and a willing partner for its dispersal (Kistler *et al*., 2015). Rind fragments and seeds recovered from archaeological sites in Arkansas, Missouri, and Kentucky indicate that between 5000 and 3000 years ago, these gourds, as cared for by people, underwent a significant transformation (Watson and Yarnell, 1966; Watson and Yarnell, 1969; Kay *et al*., 1980; Cowan, 1997; Fritz, 1997), “vegetabling” (Goldman, 2025), changing from gourds into squash. The fruits lost their bitterness due to a recessive mutation (Paris and Brown, 2005) and, over time, gradually they and their seeds increased in size (Fritz, 1997; Cowan, 1997; Kay *et al*., 1980). Likewise, over time, the fruits diversified widely in shape and topography from smooth-rinded, to lobed with sparse warts, and to heavily warted (Watson and Yarnell, 1969; Cowan, 1997).

Of the modern *Cucurbita pepo* subsp. *ovifera* squash, only the Acorn Group is cultivated for consumption of its mature fruits and, almost exclusively, only it retains the ancestral single-flowering allele (**Figure 8a**). In cucurbits, the first fruits to develop on the plant inhibit further vegetative growth, flowering, and fruit development, with inhibition beginning to take effect at 6 days past anthesis and lasting for at least 10 days (Stephenson *et al*., 1988). However, continual removal of young fruits allows the plant to sustain growth and flowering, and to produce more fruits (El-Keblawy and Lovett-Doust, 1996). Cucurbit plants produce far more flowers than necessary for maximal ripe fruit production and therefore the multiple-flowering trait would have little or no value for increased production of ripe Acorn squash.

Unlike Acorn squash, the other three modern edible-fruited morphotypes of *C. pepo* subsp. *ovifera*, Crookneck, Scallop, and Straightneck squash, are grown exclusively for the culinary use of their young fruits; indeed, their mature fruits are inedible, being toughly fibrous with hard, lignified rinds (Paris, 2000). Their fruits are picked off for eating when they are quite young, ≤ 5 days past anthesis, allowing the plants to sustain growth and flowering, and continue developing of additional fruits. The same is true of their counterparts in subsp. *pepo*, the Cocozelle and Zucchini Groups (El-Keblawy and Lovett-Doust, 1996). The plants of the three summer squash Groups of subsp. *ovifera*, though, are multiple-flowering, displaying a significantly higher number of flowers per leaf axil, and carrying the mutation at InDel#10 (**Figure 8a**, **Supplementary Table S9**). Evidently, the recessive *mf* mutation in the TCP transcription factor gene was selected long after the domestication of *Cucurbita pepo* subsp. *ovifera* var. *ozarkana*, during its subsequent evolution under cultivation as subsp. *ovifera* var. *ovifera*. During the 1500-year interval between 3000 and 1500 years ago, squash became more widely cultivated and were grown primarily for consumption of the fleshy fruits (Cowan, 1997).

The Crookneck and Scallop morphotypes are clearly contrasting extremes in summer squash of subsp. *ovifera* by the lengthening of the fruits of the former and flattening of the fruits of the latter. Interestingly, though, these two Groups are less dissimilar to one another on the basis of DNA-sequence polymorphisms than they are to the Acorn Group (Gong *et al*., 2012; Paris *et al*., 2015). This suggests that the *mf* mutation in the TCP transcription factor gene was selected by people after *Cucurbita pepo* began to be cultivated primarily for consumption of its young fruits but before the differentiation into the distinct Crookneck and Scallop Groups. Apparently, this differentiation had already occurred by 500 CE and, as Cowan (1997) observed, the archaeological remains from eastern Kentucky indicate that the fruits of that time were similar to 19^th-^ and early 20^th-^century squash grown by Native Americans.

### The quantitative nature of axillary flowering in Cucurbita pepo and utilization of allelic variation in Cpmf for breeding

Through ∼12 years of genetic and breeding effort, we were able to Mendelize the multiple-flowering trait by creation of near-isogenic lines (Paris and Hanan, 2010) (**Supplementary Figure S1**), facilitating the current positional cloning of the *Cpmf* gene. On the other hand, we showed that axillary flowering across *Cucurbita pepo* diversity is a quantitative trait and that allelic variation in InDel#10 at the *Cpmf* gene explains a significant but partial proportion of the variation (**Figure 8b**). While this analysis defines *Cpmf* as a major axillary flowering gene, the remaining variation suggests that this trait is regulated by additional genes that act independently or interact with *Cpmf*. In fact, we found significant differences in the level of axillary flowering between different subsp. *ovifera* accessions that share the multiple-flowering allele at *Cpmf* (e.g. O-SN-SNE vs. O-SC-GBS, **Figure 7d**). These are effective sources for future genetic dissection of this variation that can be explained by differences in the expression of *Cpmf* or by the actions and interactions of additional flowering-related genes in *C. pepo*.

Direct support for the interaction of *Cpmf* with genetic background is evident through the comparison of its allelic effect on axillary flowering, tendril development and harvestable fruit yield in the Cocozelle and Zucchini morphotypes. For the Cocozelle, we observed significantly stronger allelic effects of the *Cpmf* gene on fruit yield compared to the Zucchini background (**Figure 1e, f**), as well as overall enhanced axillary meristematic activity and stronger modification of tendril morphology. From a breeding standpoint, this observation implies that fine tuning of *Cpmf* effect will be essential to optimize its expression and maximize its added value.

The results of the recent introgression of *mf* into Cocozelle and Zucchini germplasm (subsp. *pepo*) from the Crookneck accession ‘Supersett’, which displayed one of the highest numbers of flowers per leaf axil of all the accessions (**Figure 7d**), has shown unequivocally the potential marked increase in yield afforded by the *mf/mf* genotype (Paris and Gur, 2022). Our recent additional breeding efforts have since shown that yield can be increased by 100% or more with deployment and optimizing the orchestration of *mf* in Cocozelle germplasm (Paris and Gur, unpublished). The ability of multiple flowering to increase yield in summer squash is reminiscent of the deployment of the “multi-pistillate” trait in cucumber (Uzcategu and Baker, 1979; Nandgaonkar and Baker, 1981; Fujieda *et al*., 1982; Anankul *et al*., 2024); however, to the best of our knowledge, a candidate or causative gene for this trait has not yet been discovered. Our results provide proof-of-concept for dissection to the causative gene level and implementation of the multiple-flowering trait in commercial breeding, and offer opportunities for precise engineering of this trait.

## MATERIALS AND METHODS

### Plant material and field experiments

#### Segregating backcross populations for mapping the mf gene

Two segregating populations were prepared. Both are ultimately derived from the initial cross, made in 1996, of a plant of *Cucurbita pepo* subsp. *pepo* Cocozelle Group ‘Striato Pugliese’, seeds obtained in 1993 from Ingegnoli, Milan, Italy, which is single-flowering, and a plant from *C. pepo* subsp. *ovifera* Crookneck Group ‘Supersett’ (SET), seeds obtained in 1990 from Harris Seeds, Rochester, New York, which is multiple-flowering. An F_1_ plant obtained from this cross was self-pollinated and several of the resulting multiple-flowering F_2_ plants were selected and crossed with two single-flowering highly inbred lines of *C. pepo* subsp. *pepo*. One of these inbreds was of Zucchini Group ‘True French’ (TRF), original seeds obtained in 1979 from Thompson & Morgan, Ipswich, U.K. The other was Accession 463, which had been derived mostly from Cocozelle Group ‘Striato d’Italia’, seeds obtained in 1989 from S.A.I.S., Cesena, Italy.

Plants derived from the cross to the Zucchini ‘True French’ were self-pollinated and several multiple-flowering F_2_ plants were backcrossed to ‘True French’. The backcross progeny were selfed, and this cycle of selection for multiple-flowering, backcrossing, and self-pollination was repeated through the F_2_ of the sixth backcross generation (**Supplementary Figure S1**). Multiple-flowering plants of that generation that were self-pollinated bred true for the multiple-flowering trait in the F_3_, and were designated as Zucchini Accession 1777 (Paris and Hanan 2010). Plants of Accession 1777 were thus nearly isogenic and resembled those of ‘True French’ except for their multiple-flowering, differentiating more than one flower bud per leaf axil.

Similarly, plants derived from the cross to the Cocozelle Accession 463 were self-pollinated and several multiple-flowering F_2_ plants were backcrossed to Accession 463. The backcross progeny were selfed, and this cycle of selection for multiple-flowering, backcrossing, and self-pollination was repeated through the F_2_ of the sixth backcross generation. Multiple-flowering plants of that generation that were self-pollinated bred true for the multiple-flowering trait in the F_3,_ and were designated as Cocozelle Accession 1951. Plants of Accession 1951 were thus nearly isogenic and resembled those of Accession 463 except for their multiple-flowering, differentiating more than one flower bud per leaf axil.

For mapping the multiple-flowering trait, we used the BC_6_F_2_ generation in both the Zucchini and Cocozelle backgrounds, as near-isogenic segregating populations. The *mf* backcrossing procedure is schematically illustrated in **Supplementary Figure S1**).

#### Phenotyping the segregating BC_6_F_2:3_ populations

Observation and scoring of the flower bud initiation was performed in the segregating BC_6_F_2:3_ populations when the plants had fully developed 15–20 internodes. Plants having several leaf axils with more than one flower bud were classified as *multiple-flowering*, whilst those having only one flower bud per leaf axil were classified as *single-flowering*.

#### Phenotyping of a core set of C. pepo accessions

Seeds of 50 accessions comprising a core collection of *Cucurbita pepo* (**Supplementary Table S1**) were sown in the field at Newe Ya‘ar on 07 April 2024. The core collection consisted of 43 edible-fruited (pumpkin and squash) and 7 inedible (gourd) accessions. The 43 pumpkin and squash accessions consisted of representatives of the eight edible-fruited cultivar-groups (Paris, 2000). The 7 gourd accessions included 3 known to be descended from collections of wild gourds and 4 in cultivation; these gourds included representatives of the two subspecies as well as one from subsp. *fraterna*, which grows wild in Mexico and is not cultivated, and one, ‘Miniature Ball’, which is offered in the seed trade but has the smallest fruits of all. ‘Miniature Ball’ holds a central position with regard to the three subspecies and is perhaps representive of their ultimate common ancestor (Gong *et al*., 2012; Paris *et al*., 2015). Accessions having bush growth habit were spaced 50 cm apart and those having vine growth habit were spaced 100 cm apart within rows, with rows spaced 200 cm apart. Ribbons were attached to the 5th, 10th, and 15th petioles on the main stem of each plant to facilitate the sequential identification of the leaf axils. Stem lengths from leaf axils 0—5 (0 being the cotyledons), 5—10, and 10—15 were measured for each plant when their respective internodes reached their full length. On each plant, from the 10th leaf through the 20th leaf, for a total of 11 per plant, the axils were observed and scored for the presence and number of branches, tendrils, leaflets, and flower buds arising directly from them, when each respective axil had enlarged and developed sufficiently to allow assessment. If a branch developed within the axil, any organs arising from it were not included in the count. From each accession, four to eight plants were scored.

### DNA extraction for whole-genome resequencing

Young leaf tissue was sampled from the parents (the Crookneck SET and the Zucchini TRF) and each plant of the BC_6_F_2_ segregating populations. DNA isolations were performed using the GenElute™ Plant Genomic Miniprep Kit (Sigma-Aldrich, St. Louis, MO, USA). DNA quality and quantification were determined using a Nanodrop ND-1000 (Nanodrop Technologies, Wilmington, DE, USA) spectrophotometer, electrophoresis on an agarose gel (1.0%), and Qubit® dsDNA BR Assay Kit (Life Technologies, Eugene, OR, USA).

### Bulk segregant analysis by sequencing (BSA-Seq)

DNA samples from young leaf tissue of 39 BC_6_F_2_ plants from the Zucchini near-isogenic population were prepared in two bulks. One bulk was derived from 24 multiple-flowering plants and the other from 15 single-flowering plants. These samples, together with DNA samples of the parents (inbreds of the Zucchini ‘True French’ and the Crookneck ‘Supersett’), were used for whole-genome resequencing (WGS) performed by Syntezza Bioscience (https://www.syntezza.com/) and Novogene (https://www.novogene.com/amea-en/). Four shotgun genomic libraries were prepared with the Hyper Library construction kit from Kapa Biosystems (Roche) with no PCR amplification. The libraries were quantitated by qPCR and sequenced on one lane for 151 cycles from each end of the fragments on a HiSeq 4000 using a HiSeq 4000 sequencing kit version1. Fastq files were generated and demultiplexed with the bcl2fastq v2.17.1.14 Conversion Software (Illumina). Average output per library was 80 million reads of 150 bp to a ∼30× coverage. All raw reads were mapped to the *Cucurbita pepo* (MU-CU-16) v4.1 reference genome (Montero-Pau *et al*., 2018) using the Burrows–Wheeler Aligner (BWA), producing analysis-ready BAM files for variant discovery with the Broad Institute’s GATK. Homozygous SNPs between the two parental alleles were extracted from the variant call format (VCF) file that was further filtered to a total depth of >20 reads per site per bulk. The read depth information for the homozygous SNPs in the multiple-flowering and single-flowering pools was obtained to calculate the SNP-index (Takagi et al., 2013). For each site, we then calculated in each bulk the ratio of the number of ‘reference’ reads to the total number of reads, which represented the SNP-index of that site. The difference between the SNP-index of two pools was calculated as ΔSNP-index. The sliding window method was used to perform the whole-genome scan and identify the trait locus confidence interval on chromosome 13.

### Seed genotyping

Genomic DNA was extracted from squash seeds in a non-destructive manner for pre-planting genotypic selection. From each seed, a small chip from the distal end (embryonic cotyledons side) of the seed was chopped off using a nail clipper. The small chips and the chopped seeds were placed in a parallel order in separate 96-well plates. The chopped-seed plates were wrapped with saran and kept in a 4°C refrigerator for as long as three months prior to sowing. DNA was extracted from the seed chips using a protocol modified from (Wang *et al*., 1993). Briefly, 75 µl of buffer A (100 mM NaOH + 2% Tween 20) were added to each well (without grinding), and the plate was then incubated for 10 minutes at 95°C. Then 75 µl of buffer B (100 mM Tris-HCl + 2 mM EDTA) were added, and the plate was mixed moderately for five minutes. Afterwards, 1–2 µl of the solution were used for PCR with the 2XPCRBIO HS Taq Mix Red (PCRBIOSYSTEMS). The annealing temperature was 56°C. Based on the genotypic results, chopped seeds were selected from the corresponding plate positions and sown, subsequently germinating into normal-appearing plants, and displayed normal germination rates.

### Fine-mapping of the *Cpmf* gene by substitution mapping

Fine mapping was performed using the BC_6_F_2_ of the Zucchini ‘True French’. Recombinants at the *mf* trait interval on chromosome 13 were scanned on the BC_6_F_2_ seeds using flanking InDel markers (**Supplementary Table S2**). BC_6_F_2_ recombinants were grown, phenotyped for their flowering pattern and self-pollinated. Recombinants that were homozygous for the multiple-flowering Crookneck parent (SET) allele on one side and heterozygous on the other side of the chromosome 13 interval were informative for mapping at this generation, while recombinants that were homozygous to the single-flowering Zucchini parent (TRF) allele on one side and heterozygous on the other side were not informative for mapping as BC_6_F_2_ progenies. All the recombinants that were used in this project for fine-mapping were phenotyped also through progeny testing at the BC_6_F_3_ generation, with 6–12 plants per family. DNA extracted from the BC_6_F_2_ recombinants was used for detailed genotyping with InDel markers at the trait interval (**Supplementary Table S2**).

### InDel marker development

For fine-mapping, InDel markers at the *mf* interval were identified based on the comparison between the parental (TRF and SET) sequences aligned to the *Cucurbita pepo* reference genome (MU-CU-16 v4.1) (Montero-Pau *et al*., 2018). Raw genomic sequences were reviewed in the target region using the Integrative Genome Viewer (IGV) (Robinson et al., 2011) and only InDels that displayed clean presence/absence of sequence reads were analyzed further. Primers were designed flanking InDel intervals and were first tested on the parents and F_1_s of the populations. PCR products were run on 2.5% agarose gel. Validated InDels, showing clear co-dominant polymorphisms, were used to genotype the BC_6_F_2_ and BC_6_F_3_ recombinants.

### Gene expression analyses

#### Sampling and mRNA extraction

Plants from the Cocozelle and Zucchini BC_6_F_2_ populations were genotyped and ∼ 10 plants were selected from each genotypic group (*Mf/Mf* and *mf/mf*) of each of the two populations, and were grown in the open field for tissue sampling. For each RNA sample, bulk tissue from two plants was used. Three biological replications were sampled for each tissue in each genotypic group. Tissue samples are illustrated in **Figure 5d**. RNA for *3’ RNA-Seq* and for Real-Time quantitative PCR (RTqPCR) was prepared with the Plant/Fungi total RNA purification kit (NORGEN).

#### 3’-mRNA-Seq

RNA-seq libraries were prepared at the Crown Genomics Institute of the Nancy and Stephen Grand Israel National Center for Personalized Medicine, Weizmann Institute of Science, Rehovot, Israel. A bulk adaptation of the MARS-Seq protocol (Jaitin *et al*., 2014; Keren-Shaul *et al*., 2019) was used to generate RNA-Seq libraries for expression profiling. Briefly, (30 ng of input) RNA from each sample was barcoded during reverse transcription and pooled. Following Agencourct Ampure XP beads cleanup (Beckman Coulter), the pooled samples underwent second-strand synthesis and were linearly amplified by T7 in vitro transcription. The resulting RNA was fragmented and converted into a sequencing-ready library by tagging the samples with Illumina sequences during ligation, RT, and PCR. Libraries were quantified by Qubit and TapeStation. Sequencing was done on a Nova-Seq X using 1.5 B, 100 cycles kit mode, allocating 1600M reads in total (Illumina). Differentially expressed genes (DEGs) between tissues or genotypes were defined as Log fold change >|1.5| and *P* value adjusted for multiple comparisons at < 0.05.

#### Real-time quantitative PCR (RTqPCR)

For cDNA preparation, we used the qScript cDNA Synthesis Kit (Quantbio). For the RTqPCR we used primers for a 111 bp amplicon of the Cp4.1LG13g07780 gene, with the gene *Elongation Factor 1-alpha* (*EF1a*, Cp4.1LG17g03150) as the reference Housekeeping gene for data standardization. Primers:

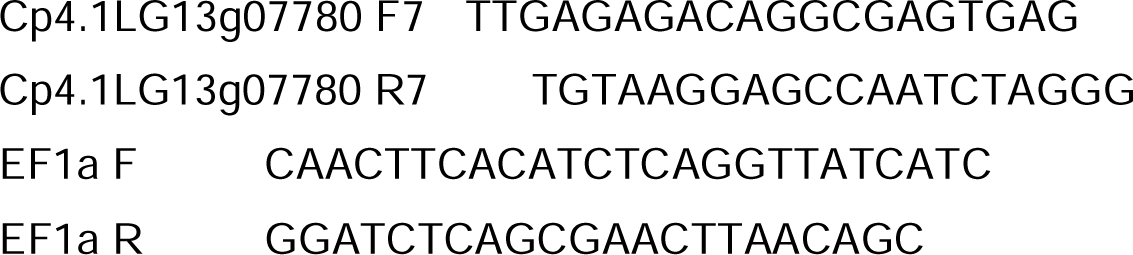

### Protein sequence comparisons and analysis of synteny

Protein sequence alignments were performed using the constraint-based alignment tool for multiple protein sequences (COBALT) (Papadopoulos and Agarwala, 2007). Synteny analyses between *Cucurbita pepo*, *Cucumis melo* and *Cucumis sativus* genomic intervals at the *Cpmf* orthologs regions were performed using the synteny viewer tool at the CuGenDBv2 database for cucurbit genomics (Yu *et al*., 2023).

### Phylogenetic analysis of *Cucurbita pepo* TCP gene family

The *Cucurbita pepo* TCP gene family was composed based on the list defined by Wang *et al*. (2025). Evolutionary analyses were conducted in MEGA12 (Kumar *et al*., 2024). The phylogenetic tree was inferred using the Neighbor-Joining method (Saitou and Nei, 1987).

### Statistical analysis

The JMP ver. 14.0.0 statistical package (SAS Institute, Cary, NC, USA) was used for all the general statistical analyses (i.e. frequency distributions, correlations, analyses of variance and mean comparisons).

## ACKNOWLEDGEMENTS

We acknowledge Zach Lippman and Anat Hendelman for helpful discussions on TCP genes and assistance with the phylogenetics of this gene family. We thank the farm team at Newe Ya‘ar for technical assistance in preparation of the field trials and for plant maintenance. Funding for this research was provided by the Israel Ministry of Agriculture Chief Scientist Grant No. 20-01-0061.

## DATA AVAILABILITY STATEMENT

The data supporting the findings of this study are available within the paper and within its supplementary materials published online. Files of genomic sequences of the parental accessions used for *Cpmf* mapping can be found at NCBI BioProject ID PRJNA1356707.

## Supplementary material

**Supplementary Figure S1:** Flow-chart describing the backcross program for introgression of *mf* into the Zucchini, ‘True-French’.

**Supplementary Figure S2:** Alignment of genomic (g) and complementary (c) DNA of *Cp4.1LG13g07780* in the parents, Crookneck ‘Supersett’ and Zucchini ‘True French’.

**Supplementary Figure S3:** Tendrils of single- and multiple-flowering Zucchini and Cocozelle near-isogenic lines.

**Supplementary Figure S4:** Fruits of the near-isogenic Zucchini and Cocozelle hybrids.

**Supplementary Figure S5**: MELO3C022091 expression is tendril-specific in melon.

**Supplementary Figure S6**: One-way analysis of total length internodes 0–15 by abbreviated pedigree.

**Supplementary Figure S7**: One-way analysis of number of male flowers per leaf axil by abbreviated pedigree.

**Supplementary Figure S8**: One-way analysis of number of female flowers per leaf axil by abbreviated pedigree.

**Supplementary Figure S9**: One-way analysis of total number of flowers per leaf axil by abbreviated pedigree.

**Supplementary Figure S10**: One-way analysis of number of tendrils per leaf axil by abbreviated pedigree.

**Supplementary Figure S11**: One-way analysis of number of branches per leaf axil by abbreviated pedigree.

**Supplementary Figure S12**: Intermediate-age fruits, 16–20 days past anthesis, of three Crookneck accessions, ‘Rugosa Friulana’ (center), ‘Early Yellow Crookneck’ (left) and ‘Yellow Summer Crookneck’ (right).

**Supplementary Figure S13**: *Cp4.1LG18g07440*, Ortholog of *CsBRC* in *Cucurbita pepo*

**Supplementary Figure S14**: Axillary flowering is independent of growth habit across a diverse core set of 50 *Cucurbita pepo* accessions.

**Supplementary Table S1:** List and description of 50 diverse *Cucurbita pepo* accessions (*Core50* panel).

**Supplementary Table S2:** Primers of InDel markers used for fine-mapping of the *Cpmf* gene on chromosome 13.

**Supplementary Table S3:** Validation of BSA-Seq results: Chr13 InDel Markers on BC_6_F_2_ tail segregants in the Zucchini and Cocozelle Groups.

**Supplementary Table S4:** List and description of 18 polymorphic sites in the *Cpmf* gene.

**Supplementary Table S5:** List of Differentially Expressed Genes (DEGs) between single-and multiple-flowering near-isogenic lines derived from the Zucchini BC_6_F_2_ in the stem at the leaf axil (SLA).

**Supplementary Table S6:** List and description of 27 auxin-related Differentially Expressed Genes (DEGs) between single- and multiple-flowering near-isogenic lines derived from the Zucchini BC_6_F_2_ in the stem at leaf axil (SLA).

**Supplementary Table S7:** List of 20 InDel markers and primers across 20 *Cucurbita pepo* chromosomes.

**Supplementary Table S8:** Genotypic data of 24 genome-wide markers across 50 diverse *Cucurbita pepo* accessions (*Core50* panel).

**Supplementary Table S9:** Genotypic data of 18 polymorphic sites in the *Cpmf* gene across 50 diverse *Cucurbita pepo* accessions (*Core50* panel).

**Supplementary Table S10:** Association analysis results of 18 polymorphic sites within the *Cpmf* gene against the mean number of flowers per leaf axil across 50 diverse *Cucurbita pepo* accessions (*Core50* panel).

## References

Aguilar-Martínez, J.A., Poza-Carrión, C. and Cubas, P. (2007) Arabidopsis Branched1 acts as an integrator of branching signals within axillary buds. Plant Cell, 19, 458–472.

Anankul, N., Sattayachiti, W., Onmanee, N., Chanmoe, S., Bundithya, W. and Kumchai, J. (2024) Genetic mapping and quantitative trait loci analysis for pistillate flowers per node and multi-pistillate flower traits in the F2 cucumber population. Breed. Sci., 74, 204–213.

Bushnell, J. (1920) The fertility and fruiting habit in Cucurbita. Proc. Am. Soc. Hortic. Sci., 17, 47–51.

Cowan, C. (1997) Evolutionary changes associated with the domestication of Cucurbita pepo. In People, Plants, and Landscapes: Studies in Paleoethnobotany. pp. 63–85.

Cubas, P., Lauter, N., Doebley, J. and Coen, E. (1999) The TCP domain: a motif found in proteins regulating plant growth and development. Plant J., 18, 215–222.

Decker-Walters, D.S., Walters, T.W., Wesley Cowan, C. and Smith, B.D. (1993) Isozymic characterization of wild populations of Cucurbita pepo. J. Ethnobiol, 13, 55–72. Available at: https://ethnobiology.org/sites/default/files/pdfs/JoE/13-1/Decker-Walters.et.al.pdf.

Dhaka, N., Bhardwaj, V., Sharma, M.K. and Sharma, R. (2017) Evolving tale of TCPs: New paradigms and old lacunae. Front. Plant Sci., 8. Available at: https://www.frontiersin.org/journals/plant-science/articles/10.3389/fpls.2017.00479.

Dixon, L.E., Greenwood, J.R., Bencivenga, S., et al. (2018) TEOSINTE BRANCHED1 regulates inflorescence architecture and development in bread wheat (Triticum aestivum). Plant Cell, 30, 563–581.

Doebley, J., Stec, A. and Hubbard, L. (1997) The evolution of apical dominance in maize. Nature, 386, 485–488.

El-Keblawy, A. and Lovett-Doust, J. (1996) Resource re-allocation following fruit removal in cucurbits: Patterns in two varieties of squash. New Phytol., 133, 583–593.

Fritz, G. (1997) A three-thousand-year-old cache of crop seed from Marble Bluff, Arkansas. In K. Gremillion, ed. People, Plants and Landscapes: Studies in Palaeoethnobotany. Tuscaloosa: University of Alabama Press, pp. 42–62.

Fujieda, K., Fujita, Y., Gunji, Y. and Takahashi, K. (1982) The inheritance of plural-pistillate flowering in cucumber. J. Japanese Soc. Hortic. Sci., 51, 172–176.

Fuller, D.Q., Denham, T., McClatchie, M. and Wu, X. (2025) Commensal domestication pathways amongst plants: Exploring segetal and ruderal crop origins. Philos. Trans. R. Soc. B Biol. Sci., 380.

Gabaldón, T. and Koonin, E. V. (2013) Functional and evolutionary implications of gene orthology. Nat. Rev. Genet., 14, 360–366.

Goldman, I.L. (2025) To vegetable: Seasons that require us. Crop Sci., 65, 1–13.

Gong, L., Paris, H.S., Nee, M.H., Stift, G., Pachner, M., Vollmann, J. and Lelley, T. (2012) Genetic relationships and evolution in Cucurbita pepo (pumpkin, squash, gourd) as revealed by simple sequence repeat polymorphisms. Theor. Appl. Genet., 124, 875–891.

Hammer, K. (1984) Das Domestikationssyndrom (1984). Kulturpflanze, 32, 11–34.

Harlan, J.R. (1992) Crops and man., American Society of Agronomy.

Heslop-Harrison, J.S. and Schwarzacher, T. (2012) Genetics and genomics of crop domestication. In A. Altman and P. M. Hasegawa, eds. Plant Biotechnology and Agriculture. San Diego: Academic Press, pp. 3–18. Available at: https://www.sciencedirect.com/science/article/pii/B9780123814661000018.

Jaitin, D.A., Kenigsberg, E., Keren-Shaul, H., et al. (2014) Massively parallel single-cell RNA-seq for marker-free decomposition of tissues into cell types. Science (80-.)., 343, 776–779. Available at: https://www.science.org/doi/abs/10.1126/science.1247651.

Kay, M., King, F.B. and Robinson, C.K. (1980) Cucurbits from Phillips Spring: New evidence and interpretations. Am. Antiq., 45, 806–822. Available at: http://www.jstor.org/stable/280151.

Keren-Shaul, H., Kenigsberg, E., Jaitin, D.A., David, E., Paul, F., Tanay, A. and Amit, I. (2019) MARS-seq2.0: an experimental and analytical pipeline for indexed sorting combined with single-cell RNA sequencing. Nat. Protoc., 14, 1841–1862. Available at: 10.1038/s41596-019-0164-4.

Kistler, L., Newsom, L.A., Ryan, T.M., Clarke, A.C., Smith, B.D. and Perry, G.H. (2015) Gourds and squashes (Cucurbita spp.) adapted to megafaunal extinction and ecological anachronism through domestication. Proc. Natl. Acad. Sci. U. S. A., 112, 15107–15112.

Kumar, S., Stecher, G., Suleski, M., Sanderford, M., Sharma, S. and Tamura, K. (2024) MEGA12: Molecular Evolutionary Genetic Analysis version 12 for adaptive and green computing. Mol. Biol. Evol., 41, msae263. Available at: 10.1093/molbev/msae263.

Loy, J.B. (2004) Morpho-physiological aspects of productivity and quality in squash and pumpkins (Cucurbita spp.). CRC. Crit. Rev. Plant Sci., 23, 337–363. Available at: 10.1080/07352680490490733.

Marcelis, L.F.M. (1993) Effect of assimilate supply on the growth of individual cucumber fruits. Physiol. Plant., 87, 313–320. Available at: 10.1111/j.1399-3054.1993.tb01736.x.

Martín-Trillo, M. and Cubas, P. (2010) TCP genes: A family snapshot ten years later. Trends Plant Sci., 15, 31–39.

McGlasson, W. and Pratt, H. (1963) Fruit-set patterns and fruit growth in cantaloupe (Cucumis melo L., var. reticulatus Naud.). Proc. Am. Soc. Hortic. Sci., 83, 495–505.

Meyer, R.S. and Purugganan, M.D. (2013) Evolution of crop species: Genetics of domestication and diversification. Nat. Rev. Genet., 14, 840–852.

Mizuno, S., Sonoda, M., Tamura, Y., Nishino, E., Suzuki, H., Sato, T. and Oizumi, T. (2015) Chiba Tendril-Less locus determines tendril organ identity in melon (Cucumis melo L.) and potentially encodes a tendril-specific TCP homolog. J. Plant Res., 128, 941–951.

Montero-Pau, J., Blanca, J., Bombarely, A., et al. (2018) De novo assembly of the zucchini genome reveals a whole-genome duplication associated with the origin of the Cucurbita genus. Plant Biotechnol. J., 16, 1161–1171.

Nandgaonkar, A.K. and Baker, L.R. (1981) Inheritance of multi-pistillate flowering habit in gynoecious pickling cucumber. J. Am. Soc. Hortic. Sci., 106, 755–757.

Nee, M. (1990) The domestication of Cucurbita (Cucurbitaceae). Econ. Bot., 44, 56–68.

Papadopoulos, J.S. and Agarwala, R. (2007) COBALT: Constraint-based alignment tool for multiple protein sequences. Bioinformatics, 23, 1073–1079.

Paris, H.S. (2018) Consumer-oriented exploitation and conservation of genetic resources of pumpkins and squash, Cucurbita. Isr. J. Plant Sci., 65, 202–221. Available at: https://brill.com/view/journals/ijps/65/3-4/article-p202_202.xml.

Paris, H.S. (2000) History of the cultivar-groups of Cucurbita pepo. In Horticultural Reviews. pp. 71–170. Available at: 10.1002/9780470650783.ch2.

Paris, H.S. (2008) Summer squash. In J. Prohens and F. Nuez, eds. In: Prohens, J., Nuez, F. (eds) Vegetables I. Handbook of Plant Breeding. New York, NY: Springer New York, pp. 351–379. Available at: 10.1007/978-0-387-30443-4_11.

Paris, H.S. and Brown, R.N. (2005) The genes of pumpkin and squash. HortScience, 40, 1620–1630.

Paris, H.S., Doron-Faigenboim, A., Reddy, U.K., Donahoo, R. and Levi, A. (2015) Genetic relationships in Cucurbita pepo (pumpkin, squash, gourd) as viewed with high frequency oligonucleotide–targeting active gene (HFO–TAG) markers. Genet. Resour. Crop Evol., 62, 1095–1111. Available at: 10.1007/s10722-015-0218-6.

Paris, H.S. and Gur, A. (2022) The multiple-flowering trait conferred by gene mf increases yield of field-grown Cocozelle and Zucchini squash. Euphytica, 218, 19. Available at: 10.1007/s10681-022-02966-5.

Paris, H.S. and Hanan, A. (2010) Single recessive gene for multiple flowering in summer squash. HortScience, 45, 1643–1644.

Paris, H.S., Lebeda, A., Křistkova, E., Andres, T.C. and Nee, M.H. (2012) Parallel evolution under domestication and phenotypic differentiation of the cultivated subspecies of cucurbita pepo (Cucurbitaceae). Econ. Bot., 66, 71–90.

Pnueli, L., Carmel-Goren, L., Hareven, D., Gutfinger, T., Alvarez, J., Ganal, M., Zamir, D. and Lifschitz, E. (1998) The SELF-PRUNING gene of tomato regulates vegetative to reproductive switching of sympodial meristems and is the ortholog of CEN and TFL1. Development, 125, 1979–1989. Available at: 10.1242/dev.125.11.1979.

Pratt, H.K., Goeschl, J.D. and Martin, F.W. (1977) Fruit growth and development, ripening, and the role of ethylene in the ‘Honey Dew’ muskmelon1. J. Am. Soc. Hortic. Sci., 102, 203–210.

Ramsay, L., Comadran, J., Druka, A., et al. (2011) INTERMEDIUM-C, a modifier of lateral spikelet fertility in barley, is an ortholog of the maize domestication gene TEOSINTE BRANCHED 1. Nat. Genet., 43, 169–172.

Rosa, J. (1924) Fruiting habit and pollination of cantaloupe. Proc. Am. Soc. Hortic. Sci., 21, 51–57.

Saitou, N. and Nei, M. (1987) The neighbor-joining method: a new method for reconstructing phylogenetic trees. Mol. Biol. Evol., 4, 406–425. Available at: 10.1093/oxfordjournals.molbev.a040454.

Schapendonk, A. and Brouwer, P. (1984) Fruit growth of cucumber in relation to assimilate supply and sink activity. Sci. Hortic. (Amsterdam)., 23, 21–33.

Shen, J., Zhang, Y., Ge, D., et al. (2019) CsBRC1 inhibits axillary bud outgrowth by directly repressing the auxin efflux carrier CsPIN3 in cucumber. Proc. Natl. Acad. Sci. U. S. A., 116, 17105–17114.

Shnaider, Y., Mitra, D., Miller, G., et al. (2018) Cucumber ovaries inhibited by dominant fruit express a dynamic developmental program, distinct from either senescence-determined or fruit-setting ovaries. Plant J., 96, 651–669.

Smith, B.D. (2006) Eastern North America as an independent center of plant domestication. Proc. Natl. Acad. Sci. U. S. A., 103, 12223–12228.

Sousa-Baena, M.S., Lohmann, L.G., Hernandes-Lopes, J. and Sinha, N.R. (2018) The molecular control of tendril development in angiosperms. New Phytol., 218, 944–958.

Stephenson, A.G., Devlin, B. and Horton, J.B. (1988) The effects of seed number and prior fruit dominance on the pattern of fruit production in Cucurbita pepo (Zucchini Squash). Ann. Bot., 62, 653–661.

Takeda, T., Suwa, Y., Suzuki, M., Kitano, H., Ueguchi-Tanaka, M., Ashikari, M., Matsuoka, M. and Ueguchi, C. (2003) The OsTB1 gene negatively regulates lateral branching in rice. Plant J., 33, 513–520.

Uzcategu, N.A. and Baker, L.R. (1979) Effects of multiple-pistillate flowering on yields of gynoecious pickling cucumbers. J. Am. Soc. Hortic. Sci., 104, 148–151.

Wang, H., Qi, M. and Cutler, A.J. (1993) A simple method of preparing plant samples for PCR. Nucleic Acids Res., 21, 4153–4154.

Wang, S., Li, W. and Jin, H. (2025) Evolution and comparison of the expression of TCP genes in the benincaseae and cucurbiteae tribes. Sci. Rep., 15, 1–16.

Wang, S., Yang, X., Xu, M., et al. (2015) A rare SNP identified a TCP transcription factor essential for tendril development in cucumber. Mol. Plant, 8, 1795–1808. Available at: 10.1016/j.molp.2015.10.005.

Watson, P.J. and Yarnell, R.A. (1966) Archaeological and paleoethnobotanical investigations in Salts Cave, Mammoth Cave National Park, Kentucky. Am. Antiq., 31, 842–849. Available at: http://www.jstor.org/stable/2694457.

Watson, P.J. and Yarnell, R.A. (1969) The prehistory of Salts Cave, Kentucky. In Reports of Investigations, No. 16. Springfield: State of Illinois. Illinois State Museum. Reports of investigations, no. 16. Springfield: [Illinois State Museum].

Yeager, A.F. (1927) Determinate growth in the tomato. J. Hered., 18, 263–265.

Yu, J., Wu, S., Sun, H., et al. (2023) CuGenDBv2: an updated database for cucurbit genomics. Nucleic Acids Res., 51, D1457–D1464. Available at: 10.1093/nar/gkac921.

Zack, C.D. and Loy, J.B. (1981) Vegetative growth of squash. Can. J. Plant Sci., 61, 673–676.

Zhang, W., Tan, L., Sun, H., et al. (2019) Natural variations at TIG1 encoding a TCP transcription factor contribute to plant architecture domestication in rice. Mol. Plant, 12, 1075–1089. Available at: 10.1016/j.molp.2019.04.005.

Zhang, Z., Deng, Y., Song, X. and Miao, M. (2015) Trehalose-6-phosphate and SNF1-related protein kinase 1 are involved in the first-fruit inhibition of cucumber. J. Plant Physiol., 177, 110–120.

Zheng, Y., Wu, S., Bai, Y., et al. (2019) Cucurbit Genomics Database (CuGenDB): A central portal for comparative and functional genomics of cucurbit crops. Nucleic Acids Res., 47, D1128–D1136.

