## Supplemental Figures for "Pleiotropic mutation in a tendril TCP gene underlies the yield-enhancing *multiple-flowering* trait in summer squash (*Cucurbita pepo*)"

**Supplementary Figure S1**: Flow-chart describing the backcross program for introgression of *mf* into the Zucchini inbred, True French.

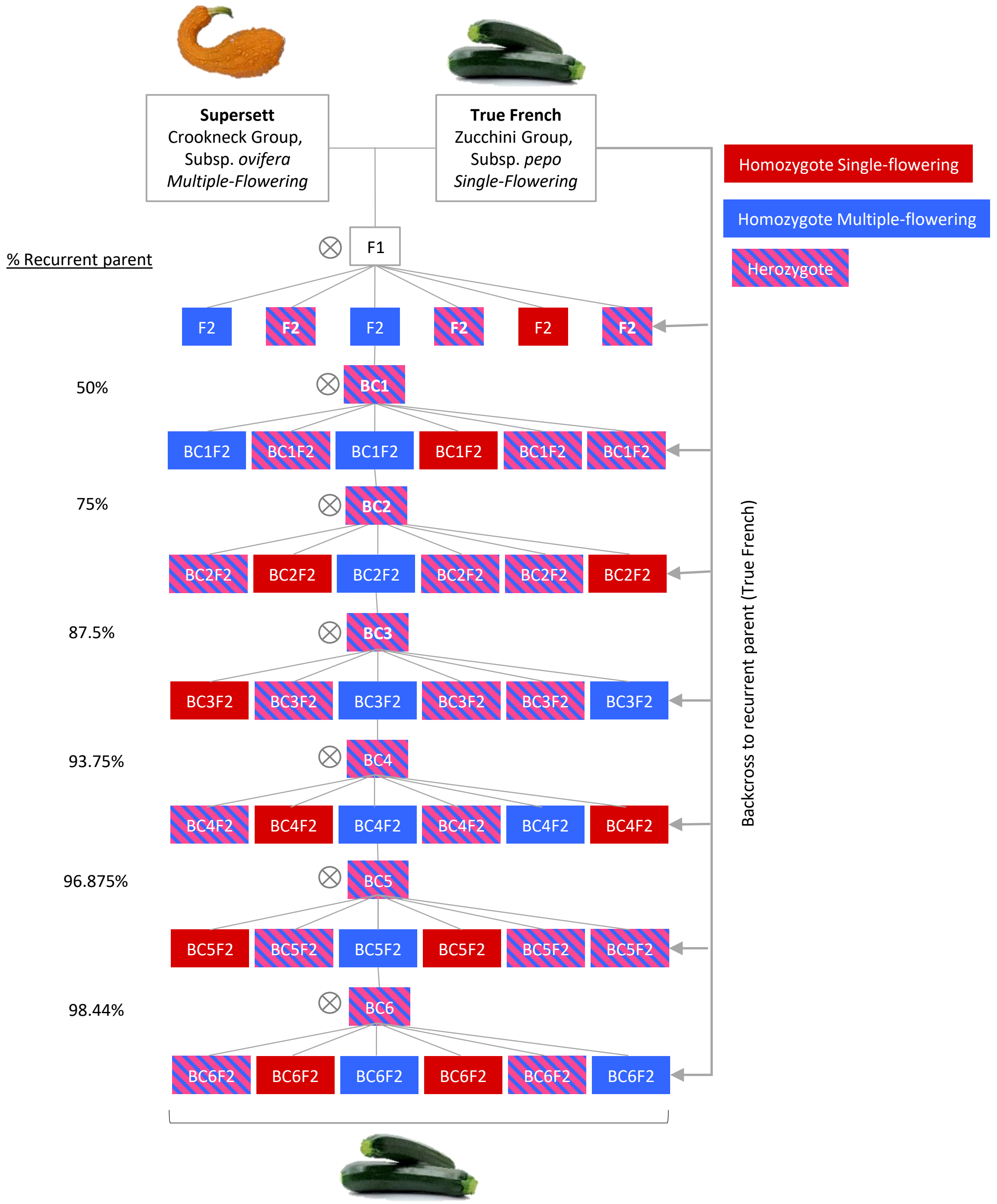

**Supplementary Figure S3:** Tendrils of Single- and multiple-flowering Zucchini and Cocozelle near-isogenic lines.

Tendrils of Single-flowering Zucchini and Cocozelle near-isogenic lines

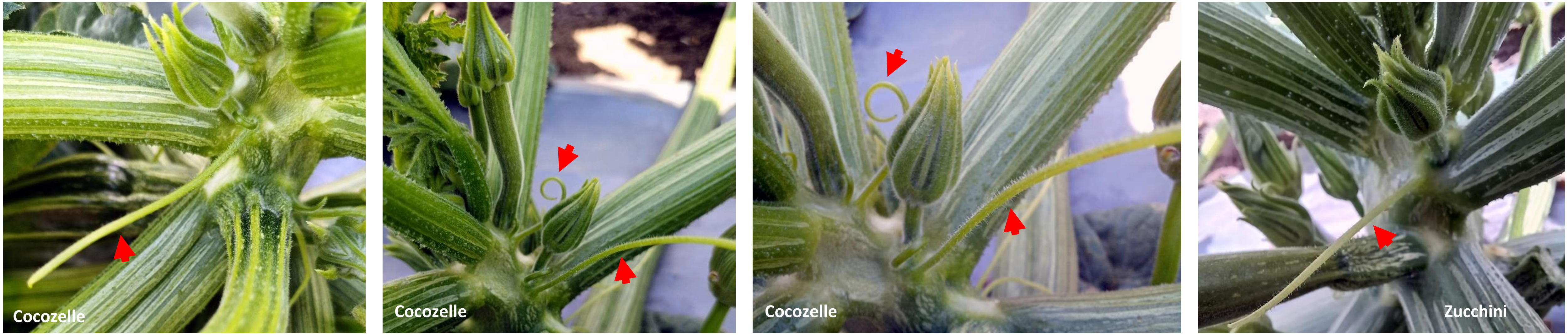


Tendrils of Multiple-flowering Zucchini and Cocozelle near-isogenic lines

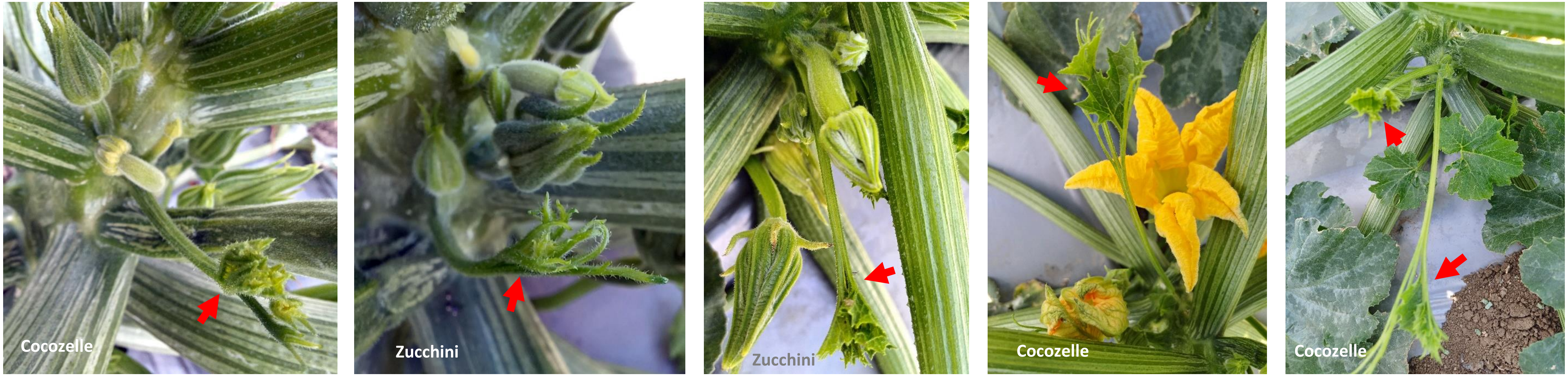

**Supplementary Figure S4:** Fruits of the near-isogenic Zucchini and Cocozelle hybrids

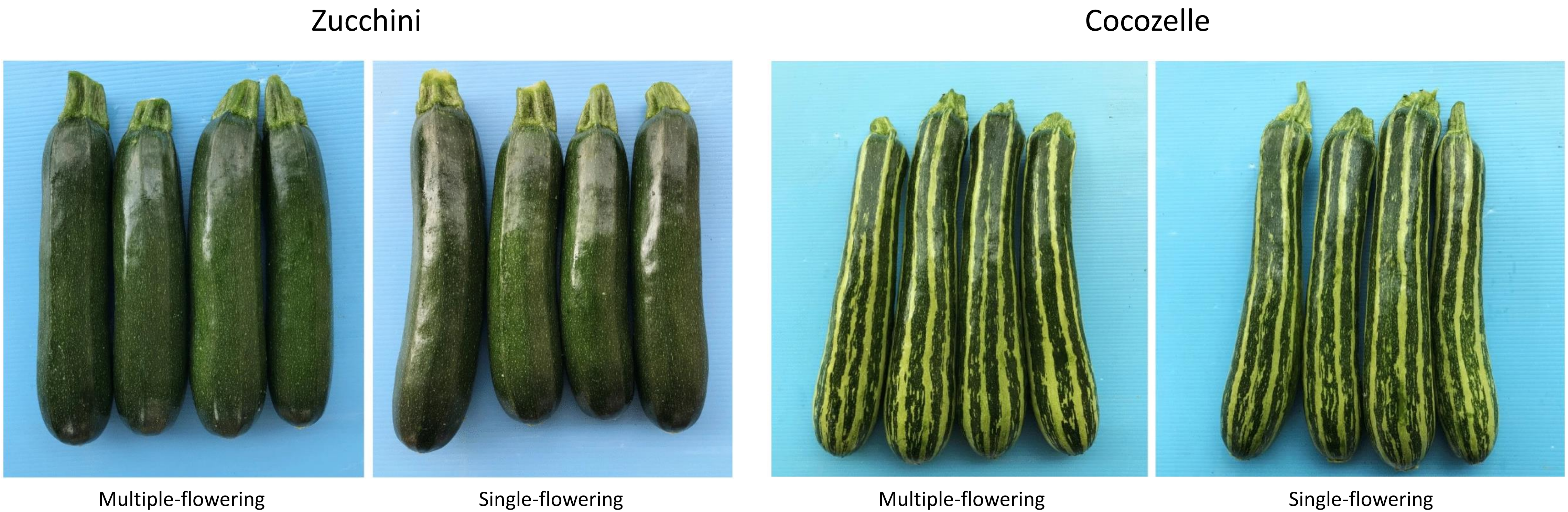


Adapted from Paris, H.S. and Gur, A. (2022) The multiple-flowering trait conferred by gene mf increases yield of field-grown Cocozelle and Zucchini squash. Euphytica, 218, 19.

**Supplementary Figure S5: MELO3C022091 expression is tendril-specific in melon**

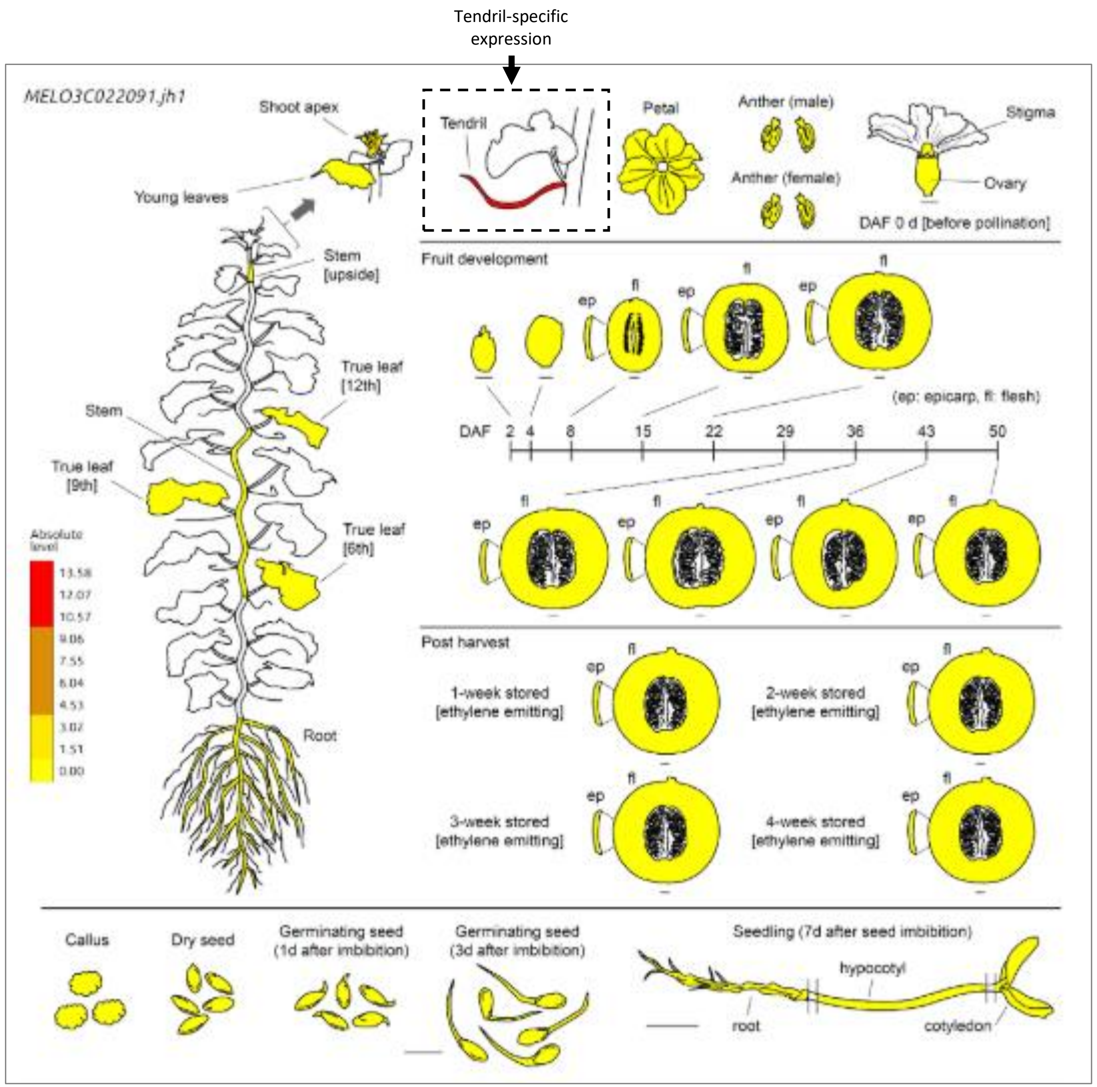


Data from Melonet-DB: https://melonet-db.dna.affrc.go.jp/ap/top

Ryoichi Yano, Satoko Nonaka, Hiroshi Ezura, Melonet-DB, a Grand RNA-Seq Gene Expression Atlas in Melon (*Cucumis melo* L.), Plant and Cell Physiology, Volume 59, Issue 1, January 2018, Page e4, https://doi.org/10.1093/pcp/pcx193

**Supplementary Figure S6**: Oneway Analysis of **Total length of Internodes 0-15** By Abbreviated pedigree

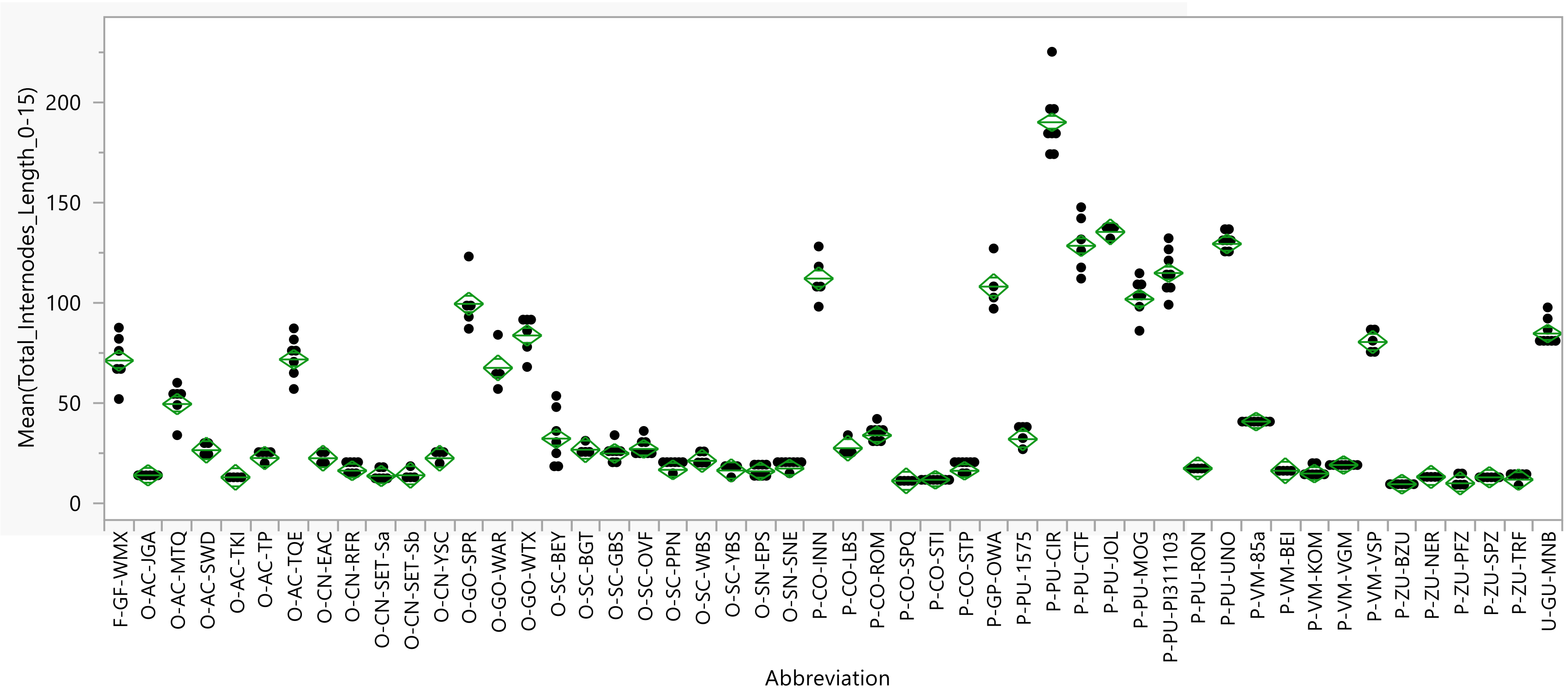


### Summary of Fit

| | |
|---|---|
| Rsquare | 0.982688 |
| Adj Rsquare | 0.979268 |
| Root Mean Square Error | 6.45644 |
| Mean of Response | 46.66443 |
| Observations (or Sum Wgts) | 298 |

### Analysis of Variance

| Source | DF | Sum of Squares | Mean Square | F Ratio | Prob > F |
|---|---|---|---|---|---|
| Abbreviation | 49 | 586834.41 | 11976.2 | 287.2984 | <.0001 * |
| Error | 248 | 10338.03 | 41.7 | | |
| C. Total | 297 | 597172.44 | | | |

### Connecting Letters Report

| Level | | Mean | Std Error |
|---|---|---|---|
| P-PU-CIR | A | 189.88 | 2.2827 |
| P-PU-JOL | B | 135.25 | 3.2282 |
| P-PU-UNO | B | 129.29 | 2.4403 |
| P-PU-CTF | B | 128.33 | 2.6358 |
| P-PU-PI311103 | C | 114.75 | 2.2827 |
| P-CO-INN | C | 112.00 | 2.8874 |
| P-GP-OWA | C D | 108.00 | 3.2282 |
| P-PU-MOG | D E | 101.71 | 2.4403 |
| O-GO-SPR | E | 99.40 | 2.8874 |
| U-GU-MNB | F | 84.63 | 2.2827 |
| O-GO-WTX | F | 83.67 | 2.6358 |
| P-VM-VSP | F | 80.40 | 2.8874 |
| O-AC-TQE | G | 71.71 | 2.4403 |
| F-GF-WMX | G | 71.17 | 2.6358 |
| O-GO-WAR | G | 67.50 | 3.2282 |
| O-AC-MTQ | H | 49.50 | 2.6358 |
| P-VM-85a | I | 40.75 | 2.2827 |
| P-CO-ROM | J | 33.88 | 2.2827 |
| O-SC-BEY | J K | 32.29 | 2.4403 |
| P-PU-1575 | J K L | 32.00 | 2.8874 |
| P-CO-LBS | J K L M | 27.50 | 3.2282 |
| O-SC-OVF | K L M | 27.13 | 2.2827 |
| O-SC-BGT | J K L M N | 26.75 | 3.2282 |
| O-AC-SWD | J K L M N | 26.50 | 3.2282 |
| O-SC-GBS | L M N | 24.86 | 2.4403 |
| O-AC-TP | M N O | 22.60 | 2.8874 |
| O-CN-EAC | M N O P | 22.50 | 3.2282 |
| O-CN-YSC | M N O P | 22.50 | 3.2282 |
| O-SC-WBS | M N O P Q | 21.20 | 2.8874 |
| P-VM-VGM | N O P Q R | 19.13 | 2.2827 |
| P-PU-RON | O P Q R S T | 17.40 | 2.8874 |
| O-SN-SNE | O P Q R S | 17.38 | 2.2827 |
| O-SC-PPN | O P Q R S T | 16.71 | 2.4403 |
| O-SC-YBS | O P Q R S T U | 16.40 | 2.8874 |
| O-CN-RFR | O P Q R S T U | 16.29 | 2.4403 |
| P-CO-STP | O P Q R S T | 16.25 | 2.2827 |
| P-VM-BEI | O P Q R S T U | 16.25 | 3.2282 |
| O-SN-EPS | O P Q R S T U | 16.00 | 2.2827 |
| P-VM-KOM | P Q R S T U | 14.88 | 2.2827 |
| O-CN-SET-Sb | P Q R S T U | 14.00 | 3.2282 |
| O-AC-JGA | Q R S T U | 14.00 | 2.6358 |
| O-CN-SET-Sa | Q R S T U | 13.67 | 2.6358 |
| P-ZU-NER | Q R S T U | 13.20 | 2.8874 |
| O-AC-TKI | Q R S T U | 13.00 | 3.2282 |
| P-ZU-SPZ | R S T U | 13.00 | 2.6358 |
| P-ZU-TRF | S T U | 11.83 | 2.6358 |
| P-CO-STI | S T U | 11.75 | 2.2827 |
| P-CO-SPQ | S T U | 11.25 | 3.2282 |
| P-ZU-PFZ | T U | 10.00 | 2.8874 |
| P-ZU-BZU | U | 9.57 | 2.4403 |

Levels not connected by same letter are significantly different.

**Supplementary Figure S7**: Oneway Analysis of **Number of male flowers** per leaf axil By abbreviated pedigree.

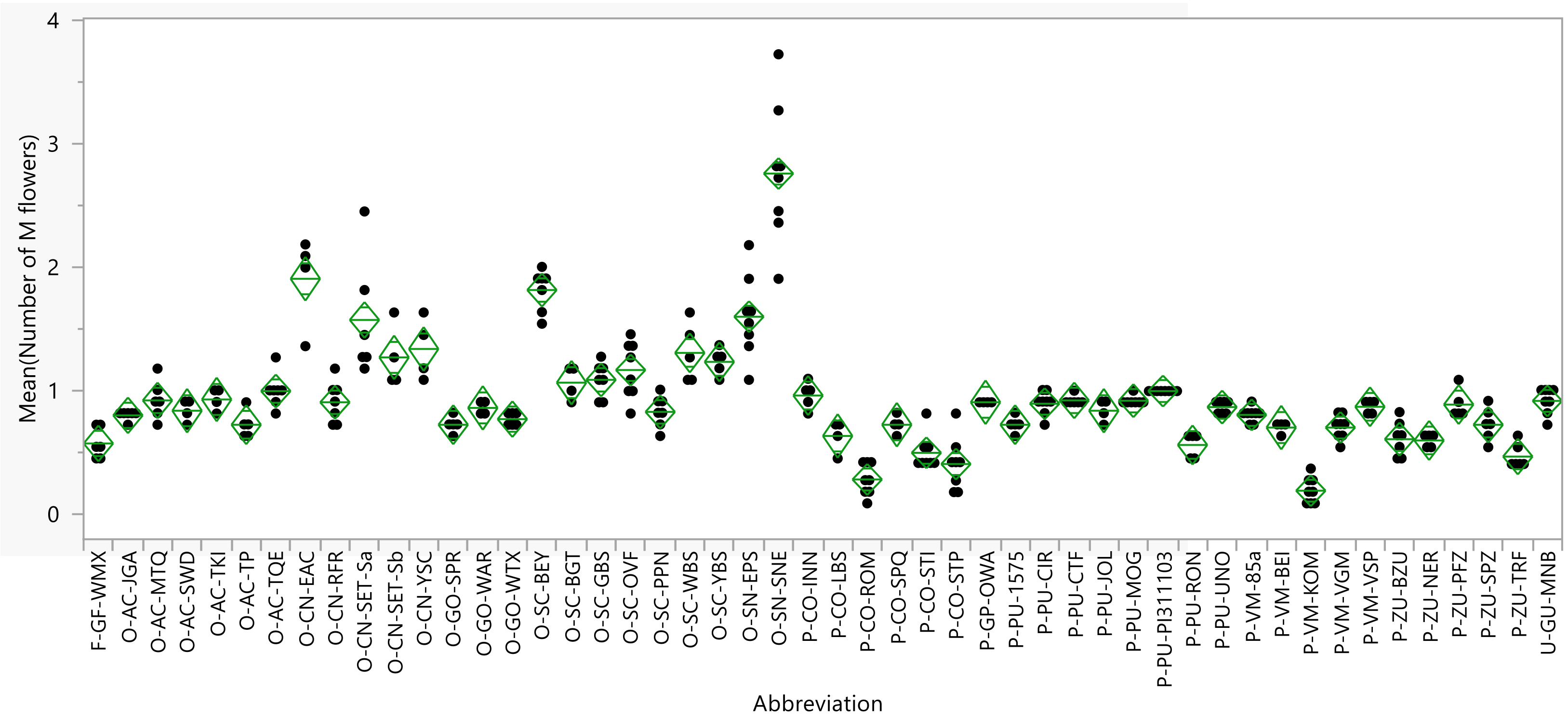


### Summary of Fit

| | |
|---|---|
| Rsquare | 0.887361 |
| Adj Rsquare | 0.865106 |
| Root Mean Square Error | 0.179972 |
| Mean of Response | 0.933801 |
| Observations (or Sum Wgts) | 298 |

### Analysis of Variance

| Source | DF | Sum of Squares | Mean Square | F Ratio | Prob > F |
|---|---|---|---|---|---|
| Abbreviation | 49 | 63.281207 | 1.29145 | 39.8721 | <.0001 * |
| Error | 248 | 8.032704 | 0.03239 | | |
| C. Total | 297 | 71.313911 | | | |

### Connecting Letters Report

| Level | | Mean | Std Error |
|---|---|---|---|
| O-SN-SNE | A | 2.7614 | 0.06363 |
| O-CN-EAC | B | 1.9091 | 0.08999 |
| O-SC-BEY | B | 1.8182 | 0.06802 |
| O-SN-EPS | C | 1.6023 | 0.06363 |
| O-CN-SET-Sa | C | 1.5758 | 0.07347 |
| O-CN-YSC | D | 1.3409 | 0.08999 |
| O-SC-WBS | D | 1.3091 | 0.08049 |
| O-CN-SET-Sb | D E | 1.2727 | 0.08999 |
| O-SC-YBS | D E | 1.2364 | 0.08049 |
| O-SC-OVF | D E F | 1.1705 | 0.06363 |
| O-SC-GBS | E F G | 1.0909 | 0.06802 |
| O-SC-BGT | E F G H | 1.0682 | 0.08999 |
| O-AC-TQE | F G H I J | 1.0000 | 0.06802 |
| P-PU-PI311103 | F G H I | 1.0000 | 0.06363 |
| P-CO-INN | G H I J K | 0.9636 | 0.08049 |
| O-AC-TKI | G H I J K L M | 0.9318 | 0.08999 |
| O-AC-MTQ | G H I J K L M | 0.9242 | 0.07347 |
| P-PU-CTF | G H I J K L M | 0.9242 | 0.07347 |
| P-PU-MOG | G H I J K L M | 0.9221 | 0.06802 |
| U-GU-MNB | G H I J K L | 0.9205 | 0.06363 |
| P-GP-OWA | G H I J K L M N | 0.9091 | 0.08999 |
| O-CN-RFR | G H I J K L M | 0.9091 | 0.06802 |
| P-PU-CIR | H I J K L M | 0.8977 | 0.06363 |
| P-ZU-PFZ | G H I J K L M N | 0.8909 | 0.08049 |
| P-VM-VSP | H I J K L M N O | 0.8727 | 0.08049 |
| P-PU-UNO | H I J K L M N | 0.8701 | 0.06802 |
| O-GO-WAR | H I J K L M N O | 0.8636 | 0.08999 |
| P-PU-JOL | H I J K L M N O | 0.8409 | 0.08999 |
| O-AC-SWD | H I J K L M N O | 0.8409 | 0.08999 |
| O-SC-PPN | I J K L M N O | 0.8312 | 0.06802 |
| P-VM-85a | K L M N O | 0.8068 | 0.06363 |
| O-AC-JGA | J K L M N O P | 0.8030 | 0.07347 |
| O-GO-WTX | K L M N O P Q | 0.7727 | 0.07347 |
| O-AC-TP | L M N O P Q | 0.7273 | 0.08049 |
| O-GO-SPR | L M N O P Q | 0.7273 | 0.08049 |
| P-PU-1575 | L M N O P Q | 0.7273 | 0.08049 |
| P-CO-SPQ | K L M N O P Q | 0.7273 | 0.08999 |
| P-ZU-SPZ | M N O P Q | 0.7273 | 0.07347 |
| P-VM-VGM | N O P Q | 0.7045 | 0.06363 |
| P-VM-BEI | L M N O P Q R | 0.7045 | 0.08999 |
| P-CO-LBS | O P Q R S | 0.6364 | 0.08999 |
| P-ZU-BZU | P Q R S | 0.6104 | 0.06802 |
| P-ZU-NER | P Q R S T | 0.6000 | 0.08049 |
| F-GF-WMX | Q R S T | 0.5758 | 0.07347 |
| P-PU-RON | Q R S T | 0.5636 | 0.08049 |
| P-CO-STI | R S T | 0.5000 | 0.06363 |
| P-ZU-TRF | S T U | 0.4697 | 0.07347 |
| P-CO-STP | T U | 0.4091 | 0.06363 |
| P-CO-ROM | U V | 0.2841 | 0.06363 |
| P-VM-KOM | V | 0.1932 | 0.06363 |

Levels not connected by same letter are significantly different.

**Supplementary Figure S8**: Oneway Analysis of **Number of female flowers** per leaf axil By abbreviated pedigree

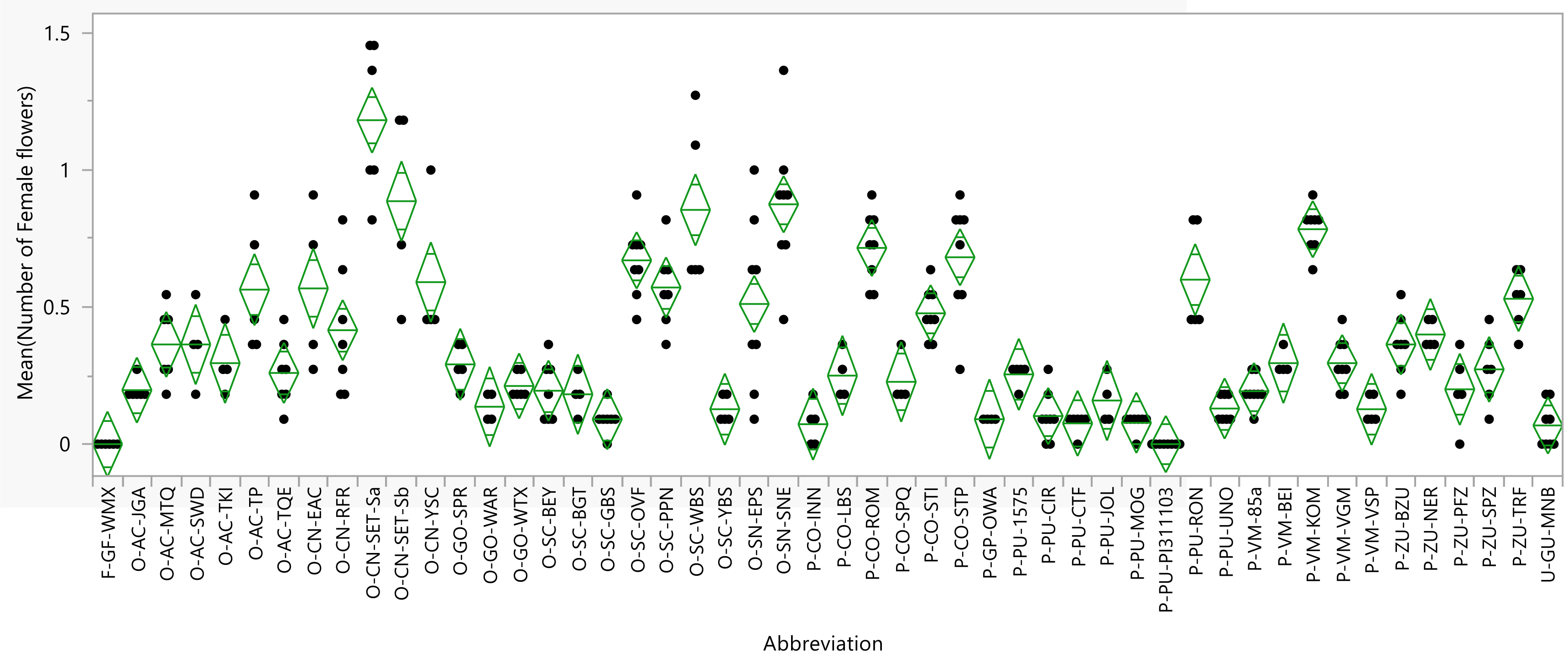


### Summary of Fit

| | |
|---|---|
| Rsquare | 0.80246 |
| Adj Rsquare | 0.76343 |
| Root Mean Square Error | 0.148235 |
| Mean of Response | 0.363636 |
| Observations (or Sum Wgts) | 298 |

### Analysis of Variance

| Source | DF | Sum of Squares | Mean Square | F Ratio | Prob > F |
|---|---|---|---|---|---|
| Abbreviation | 49 | 22.137298 | 0.451782 | 20.5601 | <.0001 * |
| Error | 248 | 5.449479 | 0.021974 | | |
| C. Total | 297 | 27.586777 | | | |

### Connecting Letters Report

| Level | | Mean | Std Error |
|---|---|---|---|
| O-CN-SET-Sa | A | 1.1818 | 0.06052 |
| O-CN-SET-Sb | B C | 0.8864 | 0.07412 |
| O-SN-SNE | B | 0.8750 | 0.05241 |
| O-SC-WBS | B C | 0.8545 | 0.06629 |
| P-VM-KOM | B C D | 0.7841 | 0.05241 |
| P-CO-ROM | C D E | 0.7159 | 0.05241 |
| P-CO-STP | D E F | 0.6818 | 0.05241 |
| O-SC-OVF | D E F | 0.6705 | 0.05241 |
| P-PU-RON | E F G | 0.6000 | 0.06629 |
| O-CN-YSC | E F G H I | 0.5909 | 0.07412 |
| O-SC-PPN | E F G H | 0.5714 | 0.05603 |
| O-CN-EAC | E F G H I J | 0.5682 | 0.07412 |
| O-AC-TP | E F G H I | 0.5636 | 0.06629 |
| P-ZU-TRF | F G H I J K | 0.5303 | 0.06052 |
| O-SN-EPS | G H I J K L | 0.5114 | 0.05241 |
| P-CO-STI | G H I J K L | 0.4773 | 0.05241 |
| O-CN-RFR | H I J K L M | 0.4156 | 0.05603 |
| P-ZU-NER | I J K L M N | 0.4000 | 0.06629 |
| O-AC-SWD | J K L M N O P Q | 0.3636 | 0.07412 |
| P-ZU-BZU | L M N O | 0.3636 | 0.05603 |
| O-AC-MTQ | K L M N O P | 0.3636 | 0.06052 |
| P-VM-VGM | M N O P Q R | 0.2955 | 0.05241 |
| O-AC-TKI | M N O P Q R S T | 0.2955 | 0.07412 |
| P-VM-BEI | M N O P Q R S T | 0.2955 | 0.07412 |
| O-GO-SPR | M N O P Q R S | 0.2909 | 0.06629 |
| P-ZU-SPZ | M N O P Q R S T | 0.2727 | 0.06052 |
| O-AC-TQE | M N O P Q R S T | 0.2597 | 0.05603 |
| P-PU-1575 | M N O P Q R S T U | 0.2545 | 0.06629 |
| P-CO-LBS | M N O P Q R S T U V | 0.2500 | 0.07412 |
| P-CO-SPQ | N O P Q R S T U V W | 0.2273 | 0.07412 |
| O-GO-WTX | O P Q R S T U V W | 0.2121 | 0.06052 |
| P-ZU-PFZ | O P Q R S T U V W | 0.2000 | 0.06629 |
| O-AC-JGA | P Q R S T U V W | 0.1970 | 0.06052 |
| O-SC-BEY | Q R S T U V W | 0.1948 | 0.05603 |
| P-VM-85a | Q R S T U V W | 0.1932 | 0.05241 |
| O-SC-BGT | O P Q R S T U V W X | 0.1818 | 0.07412 |
| P-PU-JOL | Q R S T U V W X Y | 0.1591 | 0.07412 |
| O-GO-WAR | R S T U V W X Y | 0.1364 | 0.07412 |
| P-PU-UNO | S T U V W X Y | 0.1299 | 0.05603 |
| P-VM-VSP | S T U V W X Y | 0.1273 | 0.06629 |
| O-SC-YBS | S T U V W X Y | 0.1273 | 0.06629 |
| P-PU-CIR | U V W X Y | 0.1023 | 0.05241 |
| O-SC-GBS | U V W X Y | 0.0909 | 0.05603 |
| P-GP-OWA | T U V W X Y | 0.0909 | 0.07412 |
| P-PU-MOG | V W X Y | 0.0779 | 0.05603 |
| P-PU-CTF | V W X Y | 0.0758 | 0.06052 |
| P-CO-INN | U V W X Y | 0.0727 | 0.06629 |
| U-GU-MNB | W X Y | 0.0682 | 0.05241 |
| P-PU-PI311103 | Y | 0.0000 | 0.05241 |
| F-GF-WMX | X Y | 0.0000 | 0.06052 |

Levels not connected by same letter are significantly different.

**Supplementary Figure S9**: Oneway Analysis of **Total N of flowers** per leaf axil By abbreviated pedigree

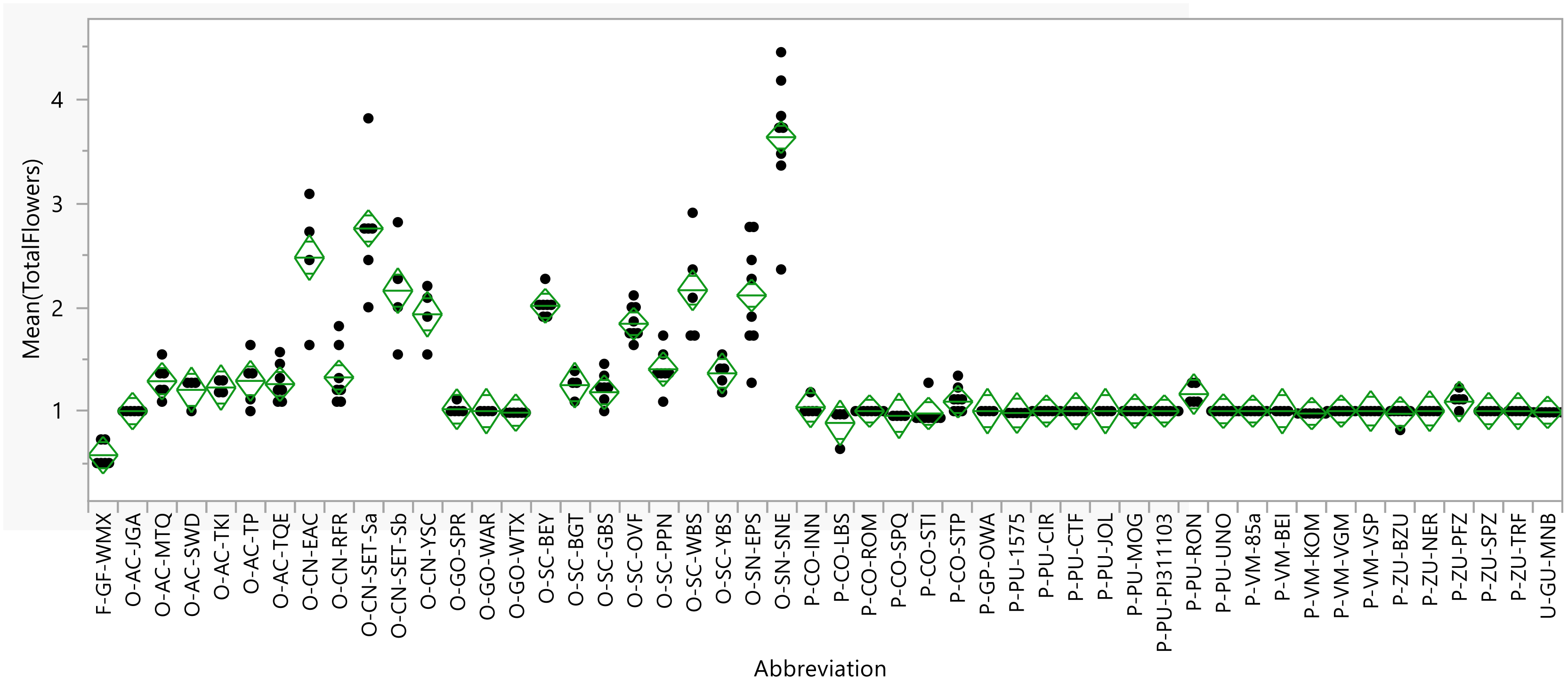


### Summary of Fit

| | |
|---|---|
| Rsquare | 0.89247 |
| Adj Rsquare | 0.871224 |
| Root Mean Square Error | 0.221592 |
| Mean of Response | 1.297437 |
| Observations (or Sum Wgts) | 298 |

### Analysis of Variance

| Source | DF | Sum of Squares | Mean Square | F Ratio | Prob > F |
|---|---|---|---|---|---|
| Abbreviation | 49 | 101.07028 | 2.06266 | 42.0069 | <.0001 * |
| Error | 248 | 12.17752 | 0.04910 | | |
| C. Total | 297 | 113.24780 | | | |

### Connecting Letters Report

| Level | | | | | | | | | | | | | Mean | Std Error |
|---|---|---|---|---|---|---|---|---|---|---|---|---|---|---|
| O-SN-SNE | A | | | | | | | | | | | | 3.6364 | 0.07834 |
| O-CN-SET-Sa | | B | | | | | | | | | | | 2.7576 | 0.09046 |
| O-CN-EAC | | B | | | | | | | | | | | 2.4773 | 0.11080 |
| O-SC-WBS | | | C | | | | | | | | | | 2.1636 | 0.09910 |
| O-CN-SET-Sb | | | C | | | | | | | | | | 2.1591 | 0.11080 |
| O-SN-EPS | | | C | | | | | | | | | | 2.1136 | 0.07834 |
| O-SC-BEY | | | C | D | | | | | | | | | 2.0130 | 0.08375 |
| O-CN-YSC | | | C | D | | | | | | | | | 1.9318 | 0.11080 |
| O-SC-OVF | | | | D | | | | | | | | | 1.8409 | 0.07834 |
| O-SC-PPN | | | | | E | | | | | | | | 1.4026 | 0.08375 |
| O-SC-YBS | | | | | E | F | | | | | | | 1.3636 | 0.09910 |
| O-CN-RFR | | | | | E | F | | | | | | | 1.3247 | 0.08375 |
| O-AC-TP | | | | | E | F | G | H | | | | | 1.2909 | 0.09910 |
| O-AC-MTQ | | | | | E | F | G | | | | | | 1.2879 | 0.09046 |
| O-AC-TQE | | | | | E | F | G | H | | | | | 1.2597 | 0.08375 |
| O-SC-BGT | | | | | E | F | G | H | I | | | | 1.2500 | 0.11080 |
| O-AC-TKI | | | | | E | F | G | H | I | J | | | 1.2273 | 0.11080 |
| O-AC-SWD | | | | | E | F | G | H | I | J | | | 1.2045 | 0.11080 |
| O-SC-GBS | | | | | E | F | G | H | I | J | | | 1.1818 | 0.08375 |
| P-PU-RON | | | | | E | F | G | H | I | J | K | | 1.1636 | 0.09910 |
| P-CO-STP | | | | | | | G | H | I | J | K | | 1.0909 | 0.07834 |
| P-ZU-PFZ | | | | | | F | G | H | I | J | K | | 1.0909 | 0.09910 |
| P-CO-INN | | | | | | | G | H | I | J | K | | 1.0364 | 0.09910 |
| O-GO-SPR | | | | | | | | H | I | J | K | | 1.0182 | 0.09910 |
| O-GO-WAR | | | | | | | | H | I | J | K | | 1.0000 | 0.11080 |
| P-VM-VSP | | | | | | | | | I | J | K | | 1.0000 | 0.09910 |
| P-CO-ROM | | | | | | | | | I | J | K | | 1.0000 | 0.07834 |
| P-GP-OWA | | | | | | | | H | I | J | K | | 1.0000 | 0.11080 |
| P-PU-CIR | | | | | | | | | I | J | K | | 1.0000 | 0.07834 |
| P-PU-CTF | | | | | | | | | I | J | K | | 1.0000 | 0.09046 |
| P-PU-JOL | | | | | | | | H | I | J | K | | 1.0000 | 0.11080 |
| P-PU-MOG | | | | | | | | | I | J | K | | 1.0000 | 0.08375 |
| P-PU-PI311103 | | | | | | | | | I | J | K | | 1.0000 | 0.07834 |
| P-PU-UNO | | | | | | | | | I | J | K | | 1.0000 | 0.08375 |
| P-VM-85a | | | | | | | | | I | J | K | | 1.0000 | 0.07834 |
| P-VM-BEI | | | | | | | | H | I | J | K | | 1.0000 | 0.11080 |
| P-VM-VGM | | | | | | | | | I | J | K | | 1.0000 | 0.07834 |
| P-ZU-NER | | | | | | | | | I | J | K | | 1.0000 | 0.09910 |
| O-AC-JGA | | | | | | | | | I | J | K | | 1.0000 | 0.09046 |
| P-ZU-SPZ | | | | | | | | | I | J | K | | 1.0000 | 0.09046 |
| P-ZU-TRF | | | | | | | | | I | J | K | | 1.0000 | 0.09046 |
| U-GU-MNB | | | | | | | | | I | J | K | | 0.9886 | 0.07834 |
| O-GO-WTX | | | | | | | | | I | J | K | | 0.9848 | 0.09046 |
| P-PU-1575 | | | | | | | | | I | J | K | | 0.9818 | 0.09910 |
| P-VM-KOM | | | | | | | | | | J | K | | 0.9773 | 0.07834 |
| P-CO-STI | | | | | | | | | | J | K | | 0.9773 | 0.07834 |
| P-ZU-BZU | | | | | | | | | | J | K | | 0.9740 | 0.08375 |
| P-CO-SPQ | | | | | | | | | I | J | K | | 0.9545 | 0.11080 |
| P-CO-LBS | | | | | | | | | | | K | | 0.8864 | 0.11080 |
| F-GF-WMX | | | | | | | | | | | | L | 0.5758 | 0.09046 |

Levels not connected by same letter are significantly different.

**Supplementary Figure S10**: Oneway Analysis of **Number of tendrils per leaf axil** By abbreviated pedigree

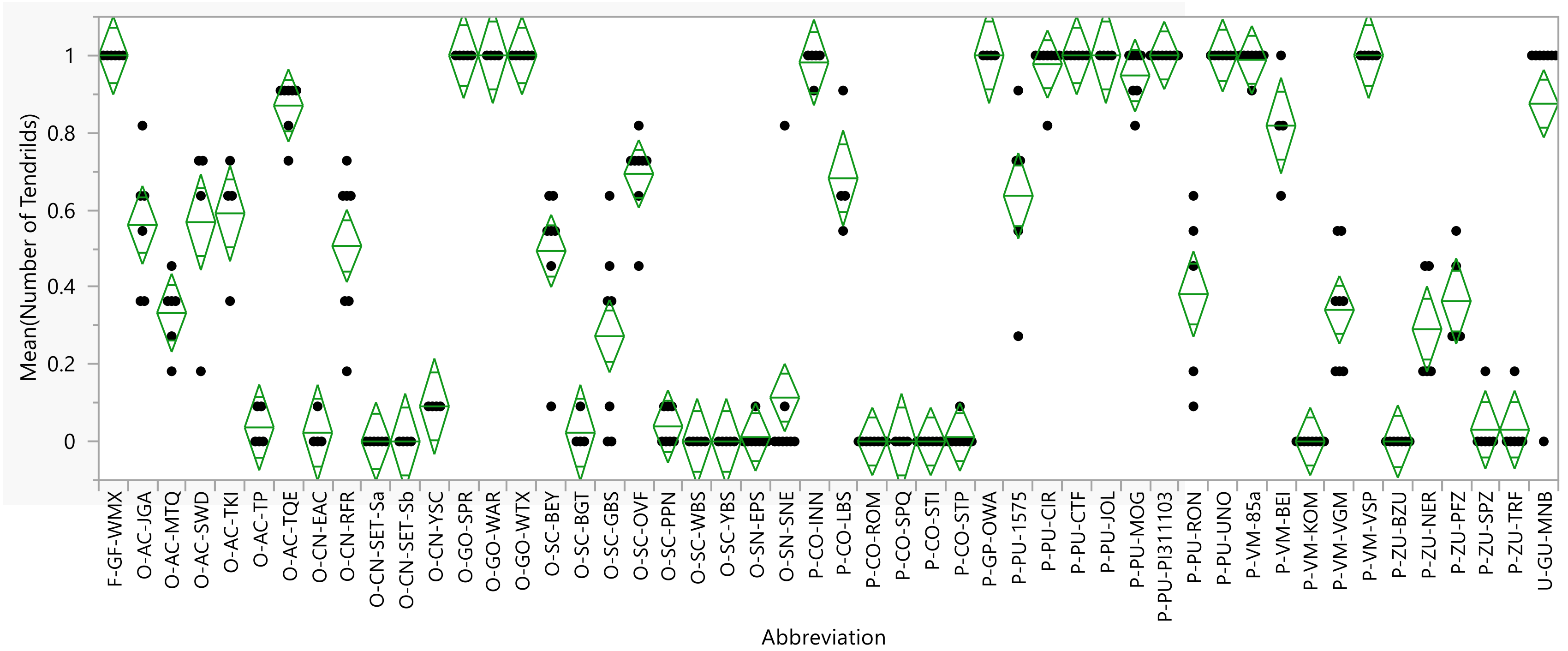


### Summary of Fit

| | |
|---|---|
| Rsquare | 0.9273 |
| Adj Rsquare | 0.912936 |
| Root Mean Square Error | 0.126228 |
| Mean of Response | 0.467663 |
| Observations (or Sum Wgts) | 298 |

### Analysis of Variance

| Source | DF | Sum of Squares | Mean Square | F Ratio | Prob > F |
|---|---|---|---|---|---|
| Abbreviation | 49 | 50.402195 | 1.02862 | 64.5572 | <.0001 * |
| Error | 248 | 3.951486 | 0.01593 | | |
| C. Total | 297 | 54.353680 | | | |

#### Connecting Letters Report

| Level | | | | | | | | | | Mean | Std Error |
|---|---|---|---|---|---|---|---|---|---|---|---|
| F-GF-WMX | A | B | | | | | | | | 1.0000 | 0.05153 |
| O-GO-WTX | A | B | | | | | | | | 1.0000 | 0.05153 |
| P-PU-UNO | A | B | | | | | | | | 1.0000 | 0.04771 |
| P-VM-VSP | A | B | | | | | | | | 1.0000 | 0.05645 |
| P-PU-PI311103 | A | | | | | | | | | 1.0000 | 0.04463 |
| O-GO-SPR | A | B | | | | | | | | 1.0000 | 0.05645 |
| P-PU-JOL | A | B | | | | | | | | 1.0000 | 0.06311 |
| P-PU-CTF | A | B | | | | | | | | 1.0000 | 0.05153 |
| P-GP-OWA | A | B | | | | | | | | 1.0000 | 0.06311 |
| O-GO-WAR | A | B | | | | | | | | 1.0000 | 0.06311 |
| P-VM-85a | A | B | | | | | | | | 0.9886 | 0.04463 |
| P-CO-INN | A | B | C | | | | | | | 0.9818 | 0.05645 |
| P-PU-CIR | A | B | | | | | | | | 0.9773 | 0.04463 |
| P-PU-MOG | A | B | C | | | | | | | 0.9481 | 0.04771 |
| U-GU-MNB | | B | C | | | | | | | 0.8750 | 0.04463 |
| O-AC-TQE | | B | C | | | | | | | 0.8701 | 0.04771 |
| P-VM-BEI | | | C | D | | | | | | 0.8182 | 0.06311 |
| O-SC-OVF | | | | D | E | | | | | 0.6932 | 0.04463 |
| P-CO-LBS | | | | D | E | | | | | 0.6818 | 0.06311 |
| P-PU-1575 | | | | | E | F | | | | 0.6364 | 0.05645 |
| O-AC-TKI | | | | | E | F | | | | 0.5909 | 0.06311 |
| O-AC-SWD | | | | | E | F | | | | 0.5682 | 0.06311 |
| O-AC-JGA | | | | | E | F | | | | 0.5606 | 0.05153 |
| O-CN-RFR | | | | | | F | G | | | 0.5065 | 0.04771 |
| O-SC-BEY | | | | | | F | G | | | 0.4935 | 0.04771 |
| P-PU-RON | | | | | | | G | H | | 0.3818 | 0.05645 |
| P-ZU-PFZ | | | | | | | G | H | | 0.3636 | 0.05645 |
| P-VM-VGM | | | | | | | | H | | 0.3409 | 0.04463 |
| O-AC-MTQ | | | | | | | | H | | 0.3333 | 0.05153 |
| P-ZU-NER | | | | | | | | H | | 0.2909 | 0.05645 |
| O-SC-GBS | | | | | | | | H | | 0.2727 | 0.04771 |
| O-SN-SNE | | | | | | | | | I | 0.1136 | 0.04463 |
| O-CN-YSC | | | | | | | | | I | 0.0909 | 0.06311 |
| O-SC-PPN | | | | | | | | | I | 0.0390 | 0.04771 |
| O-AC-TP | | | | | | | | | I | 0.0364 | 0.05645 |
| P-ZU-SPZ | | | | | | | | | I | 0.0303 | 0.05153 |
| P-ZU-TRF | | | | | | | | | I | 0.0303 | 0.05153 |
| O-CN-EAC | | | | | | | | | I | 0.0227 | 0.06311 |
| O-SC-BGT | | | | | | | | | I | 0.0227 | 0.06311 |
| P-CO-STP | | | | | | | | | I | 0.0114 | 0.04463 |
| O-SN-EPS | | | | | | | | | I | 0.0114 | 0.04463 |
| O-CN-SET-Sa | | | | | | | | | I | 0.0000 | 0.05153 |
| O-CN-SET-Sb | | | | | | | | | I | 0.0000 | 0.06311 |
| P-CO-ROM | | | | | | | | | I | 0.0000 | 0.04463 |
| O-SC-YBS | | | | | | | | | I | 0.0000 | 0.05645 |
| O-SC-WBS | | | | | | | | | I | 0.0000 | 0.05645 |
| P-CO-STI | | | | | | | | | I | 0.0000 | 0.04463 |
| P-VM-KOM | | | | | | | | | I | 0.0000 | 0.04463 |
| P-CO-SPQ | | | | | | | | | I | 0.0000 | 0.06311 |
| P-ZU-BZU | | | | | | | | | I | 0.0000 | 0.04771 |

Levels not connected by same letter are significantly different.

**Supplementary Figure S11**: Oneway Analysis of **Number of branches per leaf axil** By abbreviated pedigree

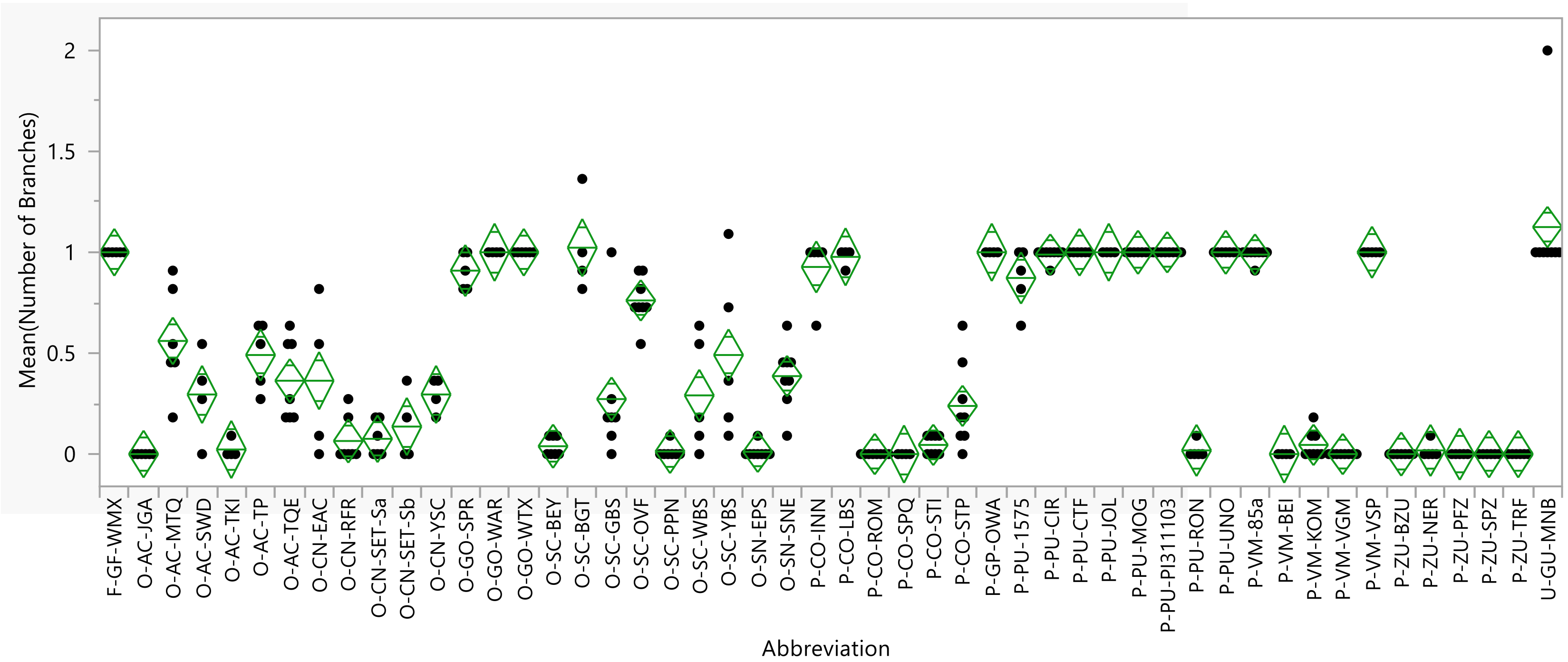


### Summary of Fit

| | |
|---|---|
| Rsquare | 0.914204 |
| Adj Rsquare | 0.897253 |
| Root Mean Square Error | 0.144776 |
| Mean of Response | 0.45424 |
| Observations (or Sum Wgts) | 298 |

### Analysis of Variance

| Source | DF | Sum of Squares | Mean Square | F Ratio | Prob > F |
|---|---|---|---|---|---|
| Abbreviation | 49 | 55.388678 | 1.13038 | 53.9305 | <.0001 * |
| Error | 248 | 5.198072 | 0.02096 | | |
| C. Total | 297 | 60.586749 | | | |

### Connecting Letters Report

| Level | | Mean | Std Error |
|---|---|---|---|
| U-GU-MNB | A | 1.1250 | 0.05119 |
| O-SC-BGT | A B | 1.0227 | 0.07239 |
| F-GF-WMX | A B | 1.0000 | 0.05910 |
| P-PU-PI311103 | A B | 1.0000 | 0.05119 |
| P-PU-MOG | A B | 1.0000 | 0.05472 |
| P-PU-JOL | A B | 1.0000 | 0.07239 |
| P-VM-VSP | A B | 1.0000 | 0.06475 |
| P-PU-CTF | A B | 1.0000 | 0.05910 |
| P-GP-OWA | A B | 1.0000 | 0.07239 |
| P-PU-UNO | A B | 1.0000 | 0.05472 |
| O-GO-WAR | A B | 1.0000 | 0.07239 |
| O-GO-WTX | A B | 1.0000 | 0.05910 |
| P-PU-CIR | A B | 0.9886 | 0.05119 |
| P-VM-85a | A B | 0.9886 | 0.05119 |
| P-CO-LBS | A B | 0.9773 | 0.07239 |
| P-CO-INN | B | 0.9273 | 0.06475 |
| O-GO-SPR | B C | 0.9091 | 0.06475 |
| P-PU-1575 | B C | 0.8727 | 0.06475 |
| O-SC-OVF | C | 0.7614 | 0.05119 |
| O-AC-MTQ | D | 0.5606 | 0.05910 |
| O-SC-YBS | D E | 0.4909 | 0.06475 |
| O-AC-TP | D E | 0.4909 | 0.06475 |
| O-SN-SNE | E F | 0.3864 | 0.05119 |
| O-CN-EAC | E F G | 0.3636 | 0.07239 |
| O-AC-TQE | E F G | 0.3636 | 0.05472 |
| O-CN-YSC | F G H | 0.2955 | 0.07239 |
| O-AC-SWD | F G H | 0.2955 | 0.07239 |
| O-SC-WBS | F G H | 0.2909 | 0.06475 |
| O-SC-GBS | F G H | 0.2727 | 0.05472 |
| P-CO-STP | G H | 0.2386 | 0.05119 |
| O-CN-SET-Sb | H I | 0.1364 | 0.07239 |
| O-CN-SET-Sa | I | 0.0758 | 0.05910 |
| O-CN-RFR | I | 0.0649 | 0.05472 |
| P-CO-STI | I | 0.0455 | 0.05119 |
| P-VM-KOM | I | 0.0455 | 0.05119 |
| O-SC-BEY | I | 0.0390 | 0.05472 |
| O-AC-TKI | I | 0.0227 | 0.07239 |
| P-PU-RON | I | 0.0182 | 0.06475 |
| P-ZU-NER | I | 0.0182 | 0.06475 |
| O-SC-PPN | I | 0.0130 | 0.05472 |
| O-SN-EPS | I | 0.0114 | 0.05119 |
| P-VM-BEI | I | 0.0000 | 0.07239 |
| P-CO-SPQ | I | 0.0000 | 0.07239 |
| P-VM-VGM | I | 0.0000 | 0.05119 |
| P-CO-ROM | I | 0.0000 | 0.05119 |
| P-ZU-BZU | I | 0.0000 | 0.05472 |
| P-ZU-PFZ | I | 0.0000 | 0.06475 |
| P-ZU-SPZ | I | 0.0000 | 0.05910 |
| P-ZU-TRF | I | 0.0000 | 0.05910 |
| O-AC-JGA | I | 0.0000 | 0.05910 |

Levels not connected by same letter are significantly different.

**Supplementary Figure S12**: 'Rugosa Friulana' (RFR) and two Crookneck accessions

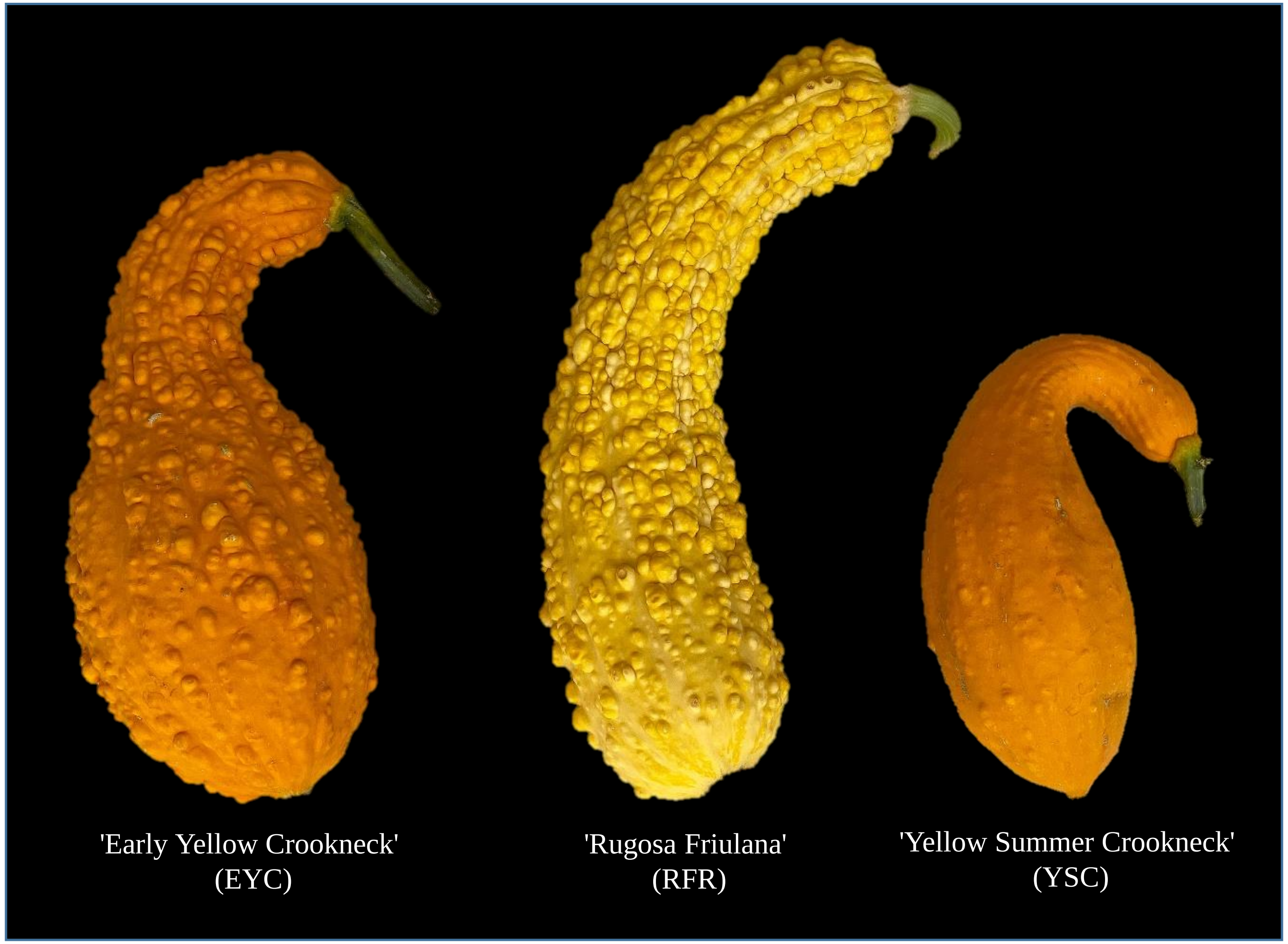

**Supplementary Figure S13**: Cp4.1LG18g07440, Ortholog of *CsBRC* in *Cucurbita pepo*

Synteny of CsBRC region with C.pepo (Zucchini) ortholog

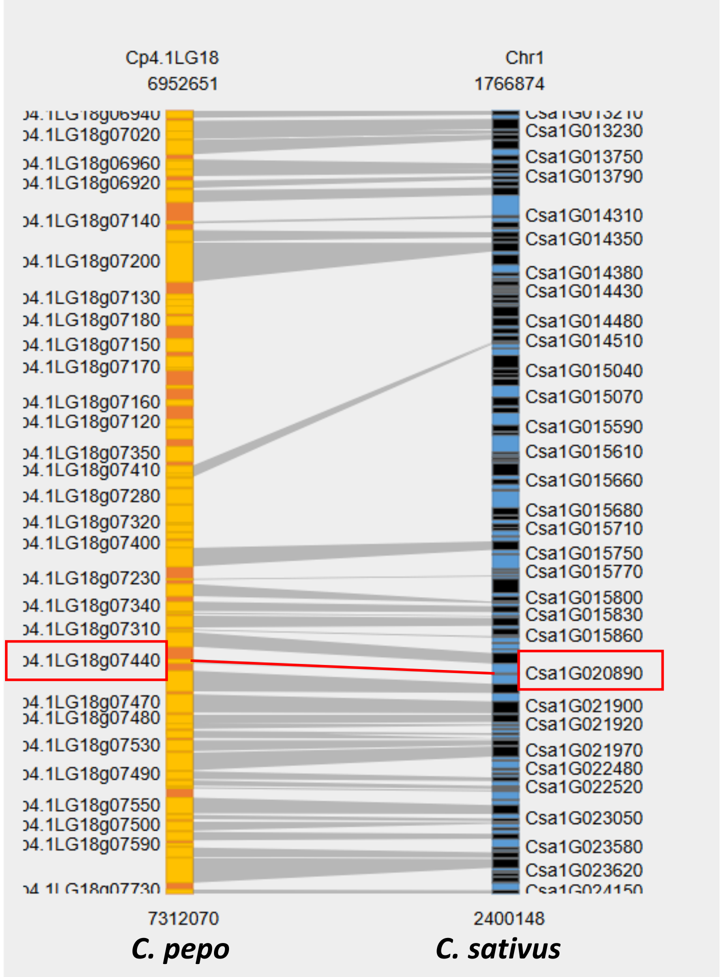


Protein BLAST on Csa1G020890 (protein) Cucumber (Chinese Long) v2

| # | Query Name | Hit Name | E-Value |
|---|---|---|---|
| 1 | unnamed protein product | Cp4.1LG18g07440.1 | 1.22227e-80 |

**Alignment**

**HSP 1**

Identity= 183/335 (54.63%) , Positive= 211/335 (62.99%) Query Matches 1 to 314 Hit Matches = 1 to 286

```
Query:    1 MFVFNSCSNNNHGNDCISFPDHSFLHFPSPFDDDGNPTNSLLLQQQSQHDIFLHHHIPLN 60
            MF+FN+       ND ISFPDHSFLHFPSPF DD  PT          Q+D  LH  I LN
Sbjct:    1 MFLFNA-------NDSISFPDHSFLHFPSPFHDD--PT-------LHQNDTLLHQ-ILLN 43

Query:   61 EPSPPPPSSTFVNALRSSET--INNDV----FHQDLVS----QRKKSSSKRDRHSKINTL 110
            EP       + FV+    S  T  I +D       HQ L+      QRK++SSKRDRHSKI+TL
Sbjct:   44 EPP-----ANFVSDFGSESTRIIVDDEHHHRVHQSLIMEQPIQRKQASSKRDRHSKIDTL 98

Query:  111 HGPRDRRMRLSLPVAKEFFGLQDMLGVDKASKTVEWLLFQARHAIKKLSKDQQSFHIDGN 170
             GPRDRRMRLSLPVA+EFFGLQDMLGVDKASKTVEWLLFQARH IKKLS    +      GN
Sbjct:   99 RGPRDRRMRLSLPVAREFFGLQDMLGVDKASKTVEWLLFQARHEIKKLSCHDRG----GN 154

Query:  171 GDTRSPSSVSDGEVVSGIIDETSTVVNNNDMISTKELEIGRKSTTKKEKRSRVGRKMPFN 230
            GD RSP S+SDGEVVSG IDE  T+VN+    I  KE       +   KKEKR RV RK  +
Sbjct:  155 GDMRSP-SISDGEVVSG-IDE--TIVNSK--IDMKE----SRKANKKEKRGRVVRKTTYY 204

Query:  231 PLTRECREKARARARARAREKQQQ-----IKGTSTTT------KLQDVSKISSPWMSSTQ 279
            PL +ECREKARARARAR  EKQ Q      +     T T       ++QDV + SS W  STQ
Sbjct:  205 PLAKECREKARARARARTIEKQLQSGKKTLSSDQTNTNKQIEGRIQDVIRTSSSW--STQ 262

Query:  280 MENNGIDEQLRTRNEGRIIMDHETTDDCLIMGRWS 314
            ++    D+QLRTRN+       ET D  ++MGRWS
Sbjct:  263 ID----DQQLRTRND-------ETDDGLMMMGRWS 286
```

**Supplementary Figure S14**: Axillary flowering is independent of growth habit across diverse core 50 *C. pepo* accessions

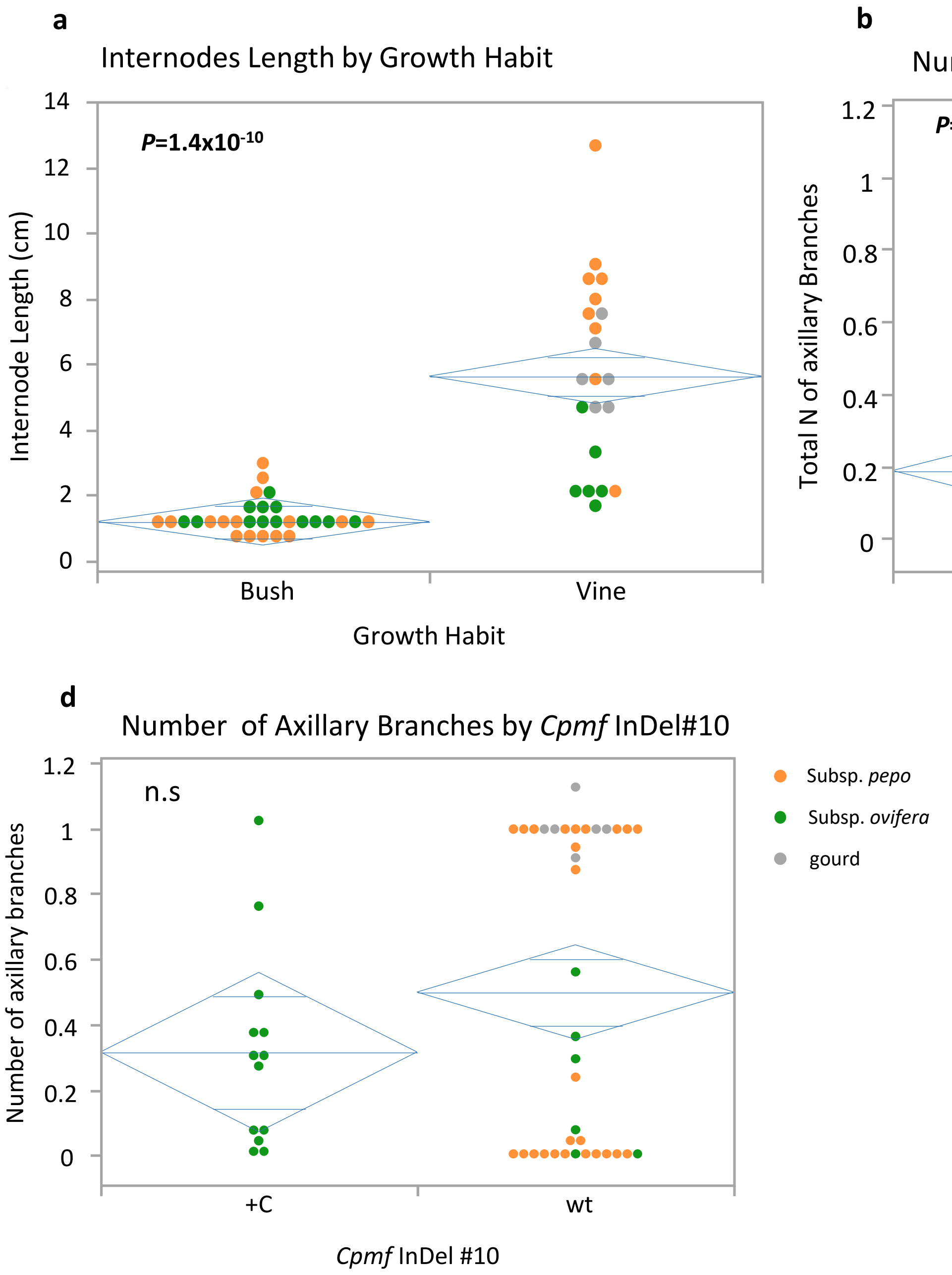

**Supplementary Figure S2**: Alignment of gDNA and cDNA of Cpmf gene (Cp4.1LG13g07780) in the Zucchini, Cocozelle and Crookneck parental accessions. (TRF: single-flowering Zucchini. 463: single-flowering, Cocozelle. SET: multiple-flowering, Crookneck).

```
                     Exon 1
                     ---------------->
gDNA_TRF             ATGTTTTCTT CCACCAACAA TCTCCACCTC TTCCCTCTTC AACACTACTT
gDNA_463             .......... ......ACAA TCTCCACCTC TTCCCTCTTC AACACTACTT
gDNA_SET             ATGTTTTCTT CCACCAACAA TCTCCGCCTC TTCCCTCTAC AACACTACTT
cDNA_463             .......... ....CAACAA TCTCCACCTC TTCCCTCTTC AACACTACTT
cDNA_SET             .......... ....CAACAA TCTCCGCCTC TTCCCTCTAC AACACTACTT
                                                SNP 1          SNP 2

gDNA_TRF             CCCTTCATCC CCCTCCCCTT ACCACCACCT CCTGCCGCCT CCTCCCCCTC
gDNA_463             CCCTTCATCC CCCTCCCCTT ACCACCACCT CCTGCCGCCT CCTCCCCCTC
gDNA_SET             CCCTTCATCC CCCTCCCCTT ACCACCACCT CCTGCCGCCT CCTCCCCCTC
cDNA_463             CCCTTCATCC CCCTCCCCTT ACCACCACCT CCTGCCGCCT CCTCCCCCTC
cDNA_SET             CCCTTCATCC CCCTCCCCTT ACCACCACCT CCTGCCGCCT CCTCCCCCTC

gDNA_TRF             CTCCGCTGCC GCCGCACCAC CCCGAGCTGA ACCCCACCAA CATCGTCTTC
gDNA_463             CTCCGCTGCC GCCGCACCAC CCCGAGCTGA ACCCCACCAA CATCGTCTTC
gDNA_SET             CTCCGCTGCC GCCGCACCAC CCCGAGCTGA ACCCCACCAA CATCGTCTTC
cDNA_463             CTCCGCTGCC GCCGCACCAC CCCGAGCTGA ACCCCACCAA CATCGTCTTC
cDNA_SET             CTCCGCTGCC GCCGCACCAC CCCGAGCTGA ACCCCACCAA CATCGTCTTC

gDNA_TRF             ATCGGCCCTC AAGATCCACC GTCGCTGCAC GGATCGGGAG GTCCATTTCT
gDNA_463             ATCGGCCCTC AAGATCCACC GTCGCTGCAC GGATCGGGAG GTCCATTTCT
gDNA_SET             ATCGGCCCTC AAGATCCACC GTCGCTGCAC GGATCGGGAG GTCCATTTCT
cDNA_463             ATCGGCCCTC AAGATCCACC GTCGCTGCAC GGATCGGGAG GTCCATTTCT
cDNA_SET             ATCGGCCCTC AAGATCCACC GTCGCTGCAC GGATCGGGAG GTCCATTTCT

gDNA_TRF             AC........ .AAGAAGAAG AAGAAGAAGA AGAAGAAGAA GCAAGGATGT
gDNA_463             AC........ .AAGAAGAAG AAGAAGAAGA AGAAGAAGAA GCAAGGATGT
gDNA_SET             ACAAGAAGAA GAAGAAGAAG AAGAAGAAGA AGAAGAAGAA GCAAGGATGT
cDNA_463             AC........ .AAGAAGAAG AAGAAGAAGA AGAAGAAGAA GCAAGGATGT
cDNA_SET             ACAAGAAGAA GAAGAAGAAG AAGAAGAAGA AGAAGAAGAA GCAAGGATGT
                         INDEL 3

gDNA_TRF             CCCAAAATGG TGGGAAGTGT TTTTCTCCTT GTGGGTTAAT TACAAAAAAA
gDNA_463             CCCAAAATGG TGGGAAGTGT TTTTCTCCTT GTGGGTTAAT TACAAAAAAA
gDNA_SET             CCCAAAATGG TGGGAAGTGT TTTTCTTCTT GTGGGTTAAT TACAAAAAAA
cDNA_463             CCCAAAATGG TGGGAAGTGT TTTTCTCCTT GTGGGTTAAT TACAAAAAAA
cDNA_SET             CCCAAAATGG TGGGAAGTGT TTTTCTTCTT GTGGGTTAAT TACAAAAAAA
                                                 SNP 4

gDNA_TRF             GGCAGTGTGA AGAAAGATCG ACATAGCAAG ATTTACACAG CTCAGGGTTT
gDNA_463             GGCAGTGTGA AGAAAGATCG ACATAGCAAG ATTTACACAG CTCAGGGTTT
gDNA_SET             GGCAGTGTGA AGAAAGATCG GCATAGCAAG ATTTACACAG CTCAGGGTTT
cDNA_463             GGCAGTGTGA AGAAAGATCG ACATAGCAAG ATTTACACAG CTCAGGGTTT
cDNA_SET             GGCAGTGTGA AGAAAGATCG GCATAGCAAG ATTTACACAG CTCAGGGTTT
                                          SNP 5

gDNA_TRF             GAGAGACAGG CGAGTGAGAT TGTCCATTGA CATTTCTAGA AAGTTCTTTG
```

```
gDNA_463        GAGAGACAGG CGAGTGAGAT TGTCCATTGA CATTTCTAGA AAGTTCTTTG
gDNA_SET        GAGAGACAGG CGAGTGAGAT TGTCCATTGA CATTTCTAGA AAGTTCTTTG
cDNA_463        GAGAGACAGG CGAGTGAGAT TGTCCATTGA CATTTCTAGA AAGTTCTTTG
cDNA_SET        GAGAGACAGG CGAGTGAGAT TGTCCATTGA CATTTCTAGA AAGTTCTTTG

gDNA_TRF        ATCTTCAAGA CATGTTGGGG TTTGACAAAG CTAGCAAAAC CCTAGATTGG
gDNA_463        ATCTTCAAGA CATGTTGGGG TTTGACAAAG CTAGCAAAAC CCTAGATTGG
gDNA_SET        ATCTTCAAGA CATGTTGGGG TTTGACAAAG CTAGCAAAAC CCTAGATTGG
cDNA_463        ATCTTCAAGA CATGTTGGGG TTTGACAAAG CTAGCAAAAC CCTAGATTGG
cDNA_SET        ATCTTCAAGA CATGTTGGGG TTTGACAAAG CTAGCAAAAC CCTAGATTGG

gDNA_TRF        CTCCTTACAA AGTCAAAAAA AGCCATTAAA GAGTTGACAA ACAACAAAAA
gDNA_463        CTCCTTACAA AGTCAAAAAA AGCCATTAAA GAGTTGACAA ACAACAAAAA
gDNA_SET        CTCCTTACAA AGTCAAAAAA AGCCATTAAA GAGTTGACAA ACAACAAAAA
cDNA_463        CTCCTTACAA AGTCAAAAAA AGCCATTAAA GAGTTGACAA ACAACAAAAA
cDNA_SET        CTCCTTACAA AGTCAAAAAA AGCCATTAAA GAGTTGACAA ACAACAAAAA

gDNA_TRF        TTTGTATAAT TTTGACTCTG AATCTGAGGC TAACCAAATA ATGTTGCCCA
gDNA_463        TTTGTATAAT TTTGACTCTG AATCTGAGGC TAACCAAATA ATGTTGCCCA
gDNA_SET        TTTGTATAAT TTTGACTCTG AATCTGAGGC TAACCAAATA ATGTTGCCCA
cDNA_463        TTTGTATAAT TTTGACTCTG AATCTGAGGC TAACCAAATA ATGTTGCCCA
cDNA_SET        TTTGTATAAT TTTGACTCTG AATCTGAGGC TAACCAAATA ATGTTGCCCA

gDNA_TRF        AATCCTCAAA CAAAACCCTT CCATTTCATG ATGTTTTGGC CAAACAATCT
gDNA_463        AATCCTCAAA CAAAACCCTT CCATTTCATG ATGTTTTGGC CAAACAATCT
gDNA_SET        AATCCTCAAA CAAAACCCTT CCATTTCATG ATGTTTTGGC CAAACAATCT
cDNA_463        AATCCTCAAA CAAAACCCTT CCATTTCATG ATGTTTTGGC CAAACAATCT
cDNA_SET        AATCCTCAAA CAAAACCCTT CCATTTCATG ATGTTTTGGC CAAACAATCT

gDNA_TRF        AGGGCAAAAG CTAGAGCAAG AGCTAGGGAA AGAACTAAGG AGAAAATCAT
gDNA_463        AGGGCAAAAG CTAGAGCAAG AGCTAGGGAA AGAACTAAGG AGAAAATCAT
gDNA_SET        AGGGCAAAAG CAAGAGCAAG AGCTAGGGAA AGAACTAAGG AGAAAATCAT
cDNA_463        AGGGCAAAAG CTAGAGCAAG AGCTAGGGAA AGAACTAAGG AGAAAATCAT
cDNA_SET        AGGGCAAAAG CAAGAGCAAG AGCTAGGGAA AGAACTAAGG AGAAAATCAT
                            SNP 7

gDNA_TRF        GAGCAAGAAT CGCCAATTCC GATCC..... .CCGCCACCG CCACCGCCTC
gDNA_463        GAGCAAGAAT CGCCAATTCC GATCC..... .CCGCCACCG CCACCGCCTC
gDNA_SET        GAGCAAGAAT CGCCAATTCC GATCCCCGCC ACCGCCACCG CCACCGCCAC
cDNA_463        GAGCAAGAAT CGCCAATTCC GATCC..... .CCGCCACCG CCACCGCCTC
cDNA_SET        GAGCAAGAAT CGCCAATTCC GATCCCCGCC ACCGCCACCG CCACCGCCAC
                                           INDEL 8                SNP 9

gDNA_TRF        CGCCTCCGCC ACCCTCCTCC TCTTCTTCTT CCATACCCGT CTTCCCAAAA
gDNA_463        CGCCTCCGCC ACCCTCCTCC TCTTCTTCTT CCATACCCGT CTTCCCAAAA
gDNA_SET        CGCCWCCGCC ACCCTCCTCC TCTTCTTCTT CCATACCCGT CTTCCCAAAA
cDNA_463        CGCCTCCGCC ACCCTCCTCC TCTTCTTCTT CCATACCCGT CTTCCCAAAA
cDNA_SET        CGCCTCCGCC ACCCTCCTCC TCTTCTTCTT CCATACCCGT CTTCCCAAAA

gDNA_TRF        GGCGGTA.CG ACCGTTGGCG GCGGAGCCAA CTTGGATCAA GAAAACAACA
gDNA_463        GGCGGTA.CG ACCGTTGGCG GCGGAGCCAA CTTGGATCAA GAAAACAACA
```

```
gDNA_SET          GGCGGTACCG ACCGTTGGCG GCGGAGCCAA CTTGGATCAA GAAAACAACA
cDNA_463          GGCGGTA.CG ACCGTTGGCG GCGGAGCCAA CTTGGATCAA GAAAACAACA
cDNA_SET          GGCGGTACCG ACCGTTGGCG GCGGAGCCAA CTTGGATCAA GAAAACAACA
                        INDEL 10

gDNA_TRF          TCATAGGGAT CATGTGGAAG CCTTCTCCTA TGATCACATC TTCAAAAGGA
gDNA_463          TCATAGGGAT CATGTGGAAG CCTTCTCCTA TGATCACATC TTCAAAAGGA
gDNA_SET          TCATAGGGAT CATGTGGAAG CCTTCTCCTA TGATCACATC TTCAAAAGGA
cDNA_463          TCATAGGGAT CATGTGGAAG CCTTCTCCTA TGATCACATC TTCAAAAGGA
cDNA_SET          TCATAGGGAT CATGTGGAAG CCTTCTCCTA TGATCACATC TTCAAAAGGA

gDNA_TRF          GAGTGTTATT ACAACAATTG CAACAATTTC TATTTTCCAC CTCCAAATTG
gDNA_463          GAGTGTTATT ACAACAATTG CAACAATTTC TATTTTCCAC CTCCAAATTG
gDNA_SET          GAGTGTTATT ACAACAATTG CAACAATTTC TATTTTCCAC CTCCAAATTG
cDNA_463          GAGTGTTATT ACAACAATTG CAACAATTTC TATTTTCCAC CTCCAAATTG
cDNA_SET          GAGTGTTATT ACAACAATTG CAACAATTTC TATTTTCCAC CTCCAAATTG

gDNA_TRF          GGACATCAAT ATTGACTCTT TTGCCAAACC ACAATCTGGG TTTTGTTCAA
gDNA_463          GGACATCAAT ATTGACTCTT TTGCCAAACC ACAATCTGGG TTTTGTTCAA
gDNA_SET          GGACATCAAT ATTGACTCTT TTGCCAAACC ACAATCTGGG TTTTGTTCAA
cDNA_463          GGACATCAAT ATTGACTCTT TTGCCAAACC ACAATCTGGG TTTTGTTCAA
cDNA_SET          GGACATCAAT ATTGACTCTT TTGCCAAACC ACAATCTGGG TTTTGTTCAA

                                             Intron 1 →
gDNA_TRF          TTGCCAACAT GGATTCTCTC CCAGGTTTTT TATTTTTATT TTTATTATTA
gDNA_463          TTGCCAACAT GGATTCTCTC CCAGGTTTTT TATTTTTATT TTTATTATTA
gDNA_SET          TTGCCAACAT GGATTCTCTC CCAGGTTTTT TATTATTATT ATTATTATTA
cDNA_463          TTGCCAACAT GGATTCTCTC CCAG...... .......... ..........
cDNA_SET          TTGCCAACAT GGATTCTCTC CCAG...... .......... ..........

gDNA_TRF          TTATTTTTAA TTATTAGGAA TCAATTTTGA AGAACATGGA TTAATGTTCT
gDNA_463          TTATTTTTAA TTATTAGGAA TCAATTTTGA AGAACATGGA TTAATGTTCT
gDNA_SET          TTTTTTTTAA TTATTAGGAA TCAATTTTGA AGAACATGGA TTAATGTTCT
cDNA_463          .......... .......... .......... .......... ..........
cDNA_SET          .......... .......... .......... .......... ..........

                                        Exon 2 →
gDNA_TRF          TGAGCTGAAA TGTTTGCAGA ATTTCAAGTT TGTGCAAAGC CATGGGATGC
gDNA_463          TGAGCTGAAA TGTTTGCAGA ATTTCAAGTT TGTGCAAAGC CATGGGATGC
gDNA_SET          TGAGCTGAAA TGTTTGCAGA ATTTCAAGTT TGTGCAAAGC CATGGGATGC
cDNA_463          .......... .........A ATTTCAAGTT TGTGCAAAGC CATGGGATGC
cDNA_SET          .......... .........A ATTTCAAGTT TGTGCAAAGC CATGGGATGC

gDNA_TRF          CTGCAGCAAC AATCAACACC TTGATCATTA A
gDNA_463          CTGCAGCAAC AATCA..... .......... .
gDNA_SET          CTGCAGCAAC AATCAACACC TTGATCATTA A
cDNA_463          CTGCAGCAAC AATCAACACC .......... .
cDNA_SET          CTGCAGCAAC AATCAAACAC .......... .
```
